# Stanniocalcin 2 (STC2) is a potent biomarker of hepatocellular carcinoma with its expression being augmented in Nrf1α-deficient cells, but diminished in Nrf2-deficient cells

**DOI:** 10.1101/2023.05.15.540796

**Authors:** Qiqi Bu, Yangxu Deng, Qing Wang, Rongzhen Deng, Shaofan Hu, Zhigang Pei, Yiguo Zhang

**Author notes:** the authors contributed equally to this work (Qiqi Bu, Yangxu Deng). Correspondence to YZ.

## Abstract

For insights into the fact that liver-specific knockout of Nrf1 leads to development of non-alcoholic steatohepatitis and spontaneous hepatoma, we previously found that loss of Nrf1α (i.e., a full-length isoform encoded by *Nfe2l1*) promotes HepG2-derived tumor growth in xenograft mice, but malgrowth of the xenograft tumor is significantly suppressed by knockout of Nrf2 (encoded by *Nfe2l2*). The mechanism underlying such marked distinctions in their pathologic phenotypes remains elusive, however, to date. Herein, we mined the transcriptome data of liver cancer from the TCGA database to establish a prognostic model of liver cancer and then calculated the predicted risk score of each cell line. The results indicated that knockout of Nrf1α significantly increased the risk score in HepG2 cells, whereas the risk score was reduced by knockout of Nrf2. Of note, stanniocalcin 2 (STC2, a biomarker of liver cancer, that is up-expressed in hepatocellular carcinoma (HCC) tissues with a reduction in the overall survival ratio of those patients) was augmented in *Nrf1αNrf2α^-/-^* cells, but diminished in Nrf2*^-/-^* cells. Thereby, it is inferable that STC2 is likely involved in mediating the distinction between *Nrf1αNrf2α^-/-^* and Nrf2*^-/-^*. Further investigation revealed that HIF1A is an upstream regulator of STC2 in caNrf2*^Δ^*^N^, rather than *Nrf1αNrf2α^-/-^*, cells, and regulation of STC2 and HIF1A in *Nrf1αNrf2α^-/-^* is determined by Nrf2, but the regulation of STC2 by Nrf2 may be independent on HIF1A. In turn, STC2 can regulate Nrf2 via the putative calcium-mediated Keap1-p62 signaling so to form a feedback regulatory loop. Such potential function of STC2 was further corroborated by a series of experiments combined with transcriptomic sequencing. The results unraveled that STC2 manifests as a dominant tumor-promoter, because the STC2-leading increases in clonogenicity of hepatoma cells and malgrowth of relevant xenograft tumor were almost completely abolished in *STC2^-/-^*cells. Together, these demonstrate that STC2 could be paved as a novel potent therapeutic target, albeit as a diagnostic marker, for hepatocellular carcinoma.

## 1. Introduction

Globally, liver cancer is a frequently-occurring malignant tumor with considerably high mortality. This is owing to the lack of clear diagnostic markers, such that it is rather difficult in gaining early diagnosis of those patients and also their prognosis is poor. Amongst all types of liver cancers, hepatocellular carcinoma (HCC) is the most common form of hematoma accounting for more than 90% [1]. This occurred closely with those increasingly unhealthy diets and lifestyles in humans, leading to non-alcoholic fatty liver disease (NAFLD), metabolic disease and obesity, which are replacing viral- and alcohol-related liver disease so as to become a core topic of HCC development [2, 3]. NAFLD is a continuum originated from the more benign course of non-alcoholic fatty liver disease (i.e., simple steatosis) and non-alcoholic steatohepatitis (NASH), which is characterized by excessive accumulation of triglycerides, with inflammation and hepatocyte damage and may culminate into liver fibrosis, cirrhosis, and even HCC [4-6]. However, the mechanisms underlying the pathogenesis of NASH and its malignant transformation to HCC remain elusive.

Coincidentally, liver-specific knockout of nuclear factor erythroid 2-related factor 1 (Nrf1, encoded by *Nfe2l1*) in mice leads to NASH and ultimately spontaneous hepatoma [7]. Further studies unraveled that Nrf1 makes a central contribution to the hepatic lipid (cholesterol) homeostasis by controlling the expression of transcriptional coactivators for the expression of metabolic enzyme genes [8-11]. Subsequently, gene expression profiling analyses revealed different pathophysiological roles of Nrf1 and Nrf2 (encoded by *Nfe2l2*) in the liver, because the former Nrf1 has limited regulation of Nrf2-target genes [12, 13], although both factors share highly evolutionary conserved homologies in the structure and function[14]. Our group found that loss of Nrf1α significantly promotes the growth of HepG2-derived xenograft tumor in nude mice, but such malgrowth of the xenograft tumor is almost completely abolished by loss of Nrf2 [15]. Collectively, such differential and even opposing phenotypes between *Nrf1αNrf2α^-/-^* and Nrf2*^-/-^* are postulated to be attributable to a hitherto unknown mechanism accounting for HCC development.

As a matter of fact, Nrf1 and Nrf2 are two principal members of the cap’n’collar (CNC) basic region-leucine zipper (bZIP) transcription factor family, which are widely expressed in a variety of tissues and cell types [16]. When they are required for biological cues, a functional heterodimer of each factor with small Mafs or other bZIP partners is formed for DNA-binding to antioxidant response elements (AREs) in their cognate gene promoter regions before such target genes are transcriptionally activated or repressed [17]. As such, ever-accumulating evidence revealed similar but distinctive roles of Nrf1 and Nrf2 in governing the transcriptional expression of proteasome, antioxidant, detoxification, metabolic and cytoprotective genes, along with those critical for maintaining cellular homeostasis. Of note, human and rodent *Nfe2l1* genes can be alternatively transcribed and further subjected to selective splicing to yield various protein isoforms with different tempo-spatial topological properties. Amongst them, the full-length Nrf1α is identified as a major player to transcriptionally regulate Nrf1-target genes [18, 19]. By contrast, Nrf2 is accepted as a master regulator of antioxidant response, but under basal conditions it is sequestered by Kelch-like ECH-associating protein 1 (Keap1) within the cytoplasm and targeted to the ubiquitin-led proteasomal degradation. Upon stimulation by oxidative stress, Nrf2 is enabled to dissociate from its inhibitor Keap1 and then translocated into the nucleus, in order to control the expression of ARE-driven genes involved in cytoprotection, differentiation, proliferation and metabolism [20]. Apart from the strong homology of between Nrf1 and Nrf2, their gene-targeting knockout experiments unraveled that they have made significantly functional differences in their pathophysiology. Knockout of Nrf1 in the mouse results in anemia due to defective erythropoiesis, leading ultimately to embryonic lethality [21], whereas Nrf2*^-/-^* mice are viable and fertile with the normal growth and development [22]. Moreover, it is of crucial significance to notice that Nrf1α is endowed as a potent tumor-repressor of hepatoma [19, 23], while Nrf2 exerts a double-edged sword’s effect on cancer development, because activation of Nrf2 is likely to inhibit NASH by ameliorating lipotoxicity, inflammation and cellular stress so to prevent liver carcinogenesis [24], but permanent oncogenic activation of Nrf2 promotes tumorigenesis and cancer malignance [15, 25].

Recently, stanniocalcin 2 (STC2) has been shown to be a tumor biomarker, which is upregulated widely in most of human cancers (e.g., hepatocellular carcinoma, esophageal carcinoma, gastric cancer, pancreatic cancer, lung cancer and prostate cancer [26-30], albeit it was originally identified as a glycoprotein hormone to regulate calcium and phosphate homeostasis. Clinical and pathological investigations also revealed that the expression abundance of STC2 is correlated with tumor progression and even prognosis of the patients. This has been exemplified by the high-level STC2 in sera from those patients with gastric cancer, which presages this pathological diagnosis and poor prognosis [31]. STC2 also seems to be correlated with the tumor size of HCC [29]; this is supported by the evidence that the overexpression of STC2 promotes cancer cell proliferation and colony formation, but conversely silencing of STC2 results in a cell-cycle delay in its G0/G1 phase. Similarly, the expression levels of STC2 in pancreatic cancer were also reported to be positively correlated with the tumor sizes, but negatively correlated with 5-year survival ratio of those patients [32]. Much to our surprise, the inducible expression of STC2 in neuronal cells was also found to be activated for response to oxidative stress and hypoxia [33]. Such hypoxia-inducible factor 1 (HIF1)-dependent expression of STC2 was further evidenced in proximal tubular epithelial cells [34]. Chromatin immunoprecipitation uncovered that HIF1A binds to the hypoxia response element (HRE) in the promoter of STC2 gene [35]. Thereby, it is inferable that STC2 can serve as a direct target of HIF1 to facilitate cell proliferation, migration and invasion under hypoxia [36, 37]. This raises an interesting question of how STC2, Nrf1 and Nrf2 together exert their essential roles in mediating the cellular response to oxidative stress, and their inter-regulatory relationship remains unknown.

To address this, we here found that STC2 can serve as a novel biomarker of hepatocellular carcinoma, with its expression being augmented in *Nrf1αNrf2α^-/-^* cells, but diminished in *^-/-^* cells. It is inferred, based on a prognostic model of liver cancer, that loss of Nrf1α led to a significant increase in the predicted risk score of hepatoma, but the risk score was reduced by loss of Nrf2. This is a full coincidence with the opposing phenotypes of their xenograft tumors in nude mice, thus implying that STC2 is likely involved in mediating such distinctions Nrf1α and Nrf2 . Further examinations revealed that upregulation of STC2 by Nrf2 in Nrf1α and caNrf2*^Δ^*^N^ cell lines, occurs via HIF1A-dependent and independent pathways, albeit Nrf2 serves as a upstream regulator of HIF1A. Conversely, STC2 regulates Nrf2 via a putative calcium-mediated Keap1-p62 signaling to form a feedback regulatory loop. The potential function of STC2 was further corroborated by a series of experiments in combination with transcriptomic sequencing. The results unraveled that like Nrf2, STC2 also manifests a dominant tumor-promoter, because STC2-promoted increases in the clonogenicity of HepG2 cells and malignant growth of its xenograft tumor were almost completely abolished in *STC2^-/-^*cells. Taken together, these demonstrate that STC2 could be paved as a novel potent therapeutic target, except as a diagnostic marker, for hepatocellular carcinoma.

## 2. Materials and methods

### 2.1. Big data mining and processing

The RNAseq data in the ‘HTSeq-Counts’ format and relative clinical information of 371 cases of hepatocellular carcinoma, along with additional 50 normal controls, were downloaded from TCGA (the Cancer Genome Atlas) website (https://portal.gdc.cancer.gov/). The normalization and differential expression analysis was performed by using the ‘DESeq2, LIMMA-voom and edgeR” packages [38]. By combination with the clinical data, both the Kaplan-Meier survival analysis and the univariate COX regression analysis were subject to establishing an eight-gene COX prognosis model for liver cancer (LIHC). The accuracy of this model was also further evaluated by using the receiver operating characteristic (ROC) curve with the concordance index.

### 2.2. Cell line culture, transfection and chemical treatment

The human hepatocellular carcinoma (HepG2) cell line was obtained originally from ATCC (Zhong Yuan Ltd., Beijing, China). Three HepG2-derived *Nrf1αNrf2α^-/-^*, Nrf2*^-/-^* (with a deletion of its transactivation domains Neh4 and Neh5) and caNrf2*^Δ^*^N^ (with a deletion of its N-terminal keap1-binding Neh2 domain by the gene-editing to yield this constitutive activation mutant) cell lines were established in our laboratory [15, 39] and identified here (Figure S1). Additional three cell lines, respectively with an insert mutant of STC2 (STC2^insC^), a knockout (KO) mutant (*STC2^-/-^*) or stably overexpressing STC2 (i.e. Lentiv-STC2), were here created from HepG2 cells and confirmed by its genomic DNA-sequencing (Figure S2). In addition, MHCC97L cell line was obtained from the Live Cancer Institute (Fudan University of China) and maintained in our laboratory.

All experimental cell lines were cultivated in DMEM (GIBCO, Life technologies) supplemented with 10% fetal bovine serum (FBS, Biological Industries, Israel), penicillin and streptomycin (100 units/mL, Solarbio, Beijing, China), in the 37°C incubator with 5% CO_2_. Two expression plasmids for human HIF1A or STC2 were constructed by cloning their cDNA sequences into the pcDNA3.1 vector. Three siRNAs (siSTC2, siNrf2 and siHIF1A) nucleotide sequences (Table S1) were synthesized for silencing their endogenous gene expression. Each of these plasmids or siRNAs was transfected into cells by incubating with Lipofectamine 3000 (Invitrogen, Carlsbad, CA, USA) for 8 h. Subsequently, the cells were allowed for a 24-h recovery from transfection in a fresh medium and then treated with the following chemicals, such as thapsigargin (TG, a microsomal Ca^2+^-ATPase inhibitor, from Sangon, A616759, Shanghai, China), CoCl_2_ (an inducer of HIF1A, from Aladdin C299372, Shanghai, China) and Oltipraz (an Nrf2 activator, that inhibits HIF1A simultaneously, from MedChemExpress HY-12519, Shanghai, China).

### 2.3. Real-time quantitative PCR

Total RNAs (1 μg) of experimental cells were extracted with the RNA simple kit (Tiangen, Beijing, China) and added to the reverse transcriptase reaction to obtain the first strand of cDNAs by using RevertAid First Strand cDNA Synthesis kit (K1622, Thermo, USA). These cDNA templates and corresponding primers (synthesized by Tsingke, Chengdu, China and listed in Table S2) were incubated with 201tμL of the real-time PCR reaction mixture including GoTaq qPCR Master Mix (Promega, USA) at 95°C for 3 min, followed by 40 cycles at 95°C for 15 s and then extending at 60°C for 30 s, in the CFX Connect Real-Time PCR Detection System (Bio-Rad, CA, USA). Therein, β-actin was used as an internal control for normalization. Subsequently, the relative mRNA expression abundances were calculated by using the 2-^△△^Ct method.

### 2.4. Western blotting with distinct antibodies

The experimental cells were collected in a lysis buffer (0.5% SDS, 0.04 mol/L DTT, pH 7.5) supplemented with the protease inhibitor EASYpacks (Roche, Germany). The lysates were diluted with 3 × loading buffer (187.5 mmol/L Tris-HCl, pH 6.8, 6% SDS, 30% Glycerol, 150 mmol/L DTT, 0.3% Bromphenol Blue), denatured for 10 min at 100°C and sonicated sufficiently. Equal amounts of protein extracts were loaded in each well of SDS-PAGE gels containing 8% or 10% polyacrylamide, and transferred to the polyvinylidene fluoride membranes (Millipore Co., Tullagreen, Ireland). The protein-loaded membranes were immunoblotted with each of the primary antibodies against STC2 (ab255610), Nrf2 (ab62352), HMOX1 (ab68477), NQO1(ab80588), *GCLM* (Ab126704) (all five antibodies purchased from Abcam), HIF1A (#36169, from Cell Signaling Technology), V5 tag (R960-25, from Thermo Fisher), Nrf1 (this specific antibody made in our own laboratory [40]), or β-actin (TA-09, from ZSGB-BIO, Beijing, China) overnight at 4°C and then the secondary antibodies [HRP-labeled goat anti-rabbit or anti-mouse IgG (H+L), ZB-2301, from ZSGB-BIO, Beijing, China] for 2 h at 37°C. The immunoblots were lastly exposed to the ECL light system and calculated by using the ImageJ software.

### 2.5. The STC2 gene-editing by CRISPR/Cas9 to yield STC2 and

The gRNA-target sequences were designed online (http://crispr.dbcls.jp/) (Table S3) and then cloned into the Cas9/Grna (puro-GFP) Vector (Wiewsolid Biotech, China). The indicated plasmids were confirmed by sequencing and co-transfected into HepG2 cells for 8 h, before the cells were allowed for a 24-h recovery from transfection in a complete medium containing 10% FBS. The transfected cells were screened with puromycin (Solarbio, Beijing, China), diluted and inoculated into 96-well cell culture plates (with a probability of one cell per well). The positively-selected monoclonal cell lines were subjected to the genomic DNA extraction and PCR amplification of the gRNA-target-adjoining sequences to identify their genotypes, called STC2^insC^ and *STC2^-/-^*, respectively.

### 2.6. STC2-overexpressing cell lines were established by lentivirus

The STC2-encoding cDNA was cloned into the pLJM1-EGFP vector to yield a STC2-expressing construct, called pLJM1-STC2, that was verified by sequencing. The pLJM1-STC2, together with the virus-packaging plasmids psPAX2 and pMD2G, was co-transfected into 293T cells for 8 h, before these cells were allowed for a 24-h recovery from transfection in a complete medium containing 10% FBS. The cells continued to culture for additional 24 h and their supernatants were collected to obtain a certain amount of lentivirus. The lentivirus titer was then evaluated, prior to an efficient infection of the STC2-expressing lentivirus into HepG2 cells. The positive monoclonal cells (called Lentiv-STC2) were selected and saved for subsequent experiments.

### 2.7. The colony formation assay

Experimental cells (750 cells/well seeded in 6-well plates) were allowed for growing for two weeks at 37 °C with 5% CO_2_. The cells were subjected to fixation by 4% paraformaldehyde before being stained with 1% crystal violet reagent (Sigma), and then the colony number was counted.

### 2.8. Analysis of cell cycle by flow cytometry

The experimental cells were collected by centrifuging at 1000 rpm for 5 min, and suspended in 300 μL of pre-cooled PBS, before being slowly added in 700 μL of absolute ethanol, gently mixed, and then incubated at 4°C overnight. The cells were re-centrifuged at 4°C and re-suspended in 100 μL of a binding buffer. The cell suspensions were incubated in the dark with 5 μL of propidium iodide (PI)-staining solution and 5 μL of annexin V-FITC at room temperature for 15 min. Additional 400 μL of binding buffer was added to the cell sample and mixed fully before analysis of the cell cycle by flow cytometer.

### 2.9. Subcutaneous tumor xenografts in nude mice

Mouse xenograft tumor models were made by subcutaneously heterotransplanting the wild type (WT) HepG2 and its derived STC2^insC^, *^-/-^* and Lentiv-STC2 cell lines into nude mice, as described [41]. Each of experimental cell lines (1×10^8^) in exponential growth phase were suspended in 0.1 mL of phosphate buffer solution, before being inoculated subcutaneously into the indicated axilla region of male nude mice (BALB/C^nu/nu^, 6 weeks, 18 g, from HFK Bioscience, Beijing, China) at a single site (n = 5 per group). The inoculated procedure into all mice was completed within 30 min, and the formation of subsequent subcutaneous tumor xenografts was observed. The tumor sizes were measured every two days until the 30^nd^ day when all mice were sacrificed and also their transplanted tumors were excised. The sizes of all xenograft tumors were calculated by a standard formula (i.e., V = ab^2^/2). Notably, all mice were maintained under standard diets and living conditions. All relevant studies were carried out on the mice (with the license No. PIL60/13167) in accordance with United Kingdom Animal (Scientific Procedures) Act (1986) and the guidelines of the Animal Care and Use Committees of Chongqing University and the Third Military Medical University, both of which were subjected to the local ethical review (in China). All relevant experimental protocols were approved by the University Laboratory Animal Welfare and Ethics Committee (with two institutional licenses SCXK-PLA-20120011 and SYXK-PLA-20120031).

### 2.10. Analysis of transcriptome sequencing

Total RNAs were extracted from *WT*, *STC2^-/-^* and Lentiv-STC2 cell lines and subjected to their transcriptome sequencing (by Beijing Genomics Institute, Shenzhen, China) on the DNBSEQ platform. After the data were filtered, those clean reads are obtained and then mapped to the relevant reference sequences of Homo sapiens’ genome (GCF_000001405.38_GRCh38.p12) by using both tools HISAT [42] and Bowtie2 [43]. The relative gene expression levels of each sample were calculated by using the RSEM method [44]. Consequently, the differentially expressed genes (DEGs) were identified, with a criteria Log_2_ fold-changes ≥1 and Q-value ≤0.05, by using the DESeq2 tool. Those DEGs were further subjected to both the Gene Ontology (GO [45]) functional enrichment analysis (including biological processes, cellular components and molecular functions) and also the Kyoto Encyclopedia of Genes and Genomes (KEGG) [46] pathway enrichment analysis.

### 2.11. Statistical analysis

All relevant data in this study were obtained from at least three independent experiments, each of which was performed in triplicates and shown as fold changes (mean1t±1tSD), before being analyzed by using the Origin8.0 tool. The statistic differences between the various experimental groups and within groups were calculated by one-way ANOVA, and the results at the value p1t< 01t.01 were considered to have significant differences.

## 3. Results

### 3.1. Involvement of STC2 in mediating the distinction between **Nrf1α**^-/-^ and *Nrf2*α^-/-^

To gain insight into distinct phenotypes of those tumor xenograft mice inoculated with *Nrf1αNrf2α^-/-^*, rather than Nrf2, hepatoma cell lines, we first analyzed the data obtained from the TCGA database by using distinct packages (Figures S3 & S4). The results of principal component analysis (PCA) of HCC and normal cases indicated in the TCGA database were shown (Figure S4A), along with a volcano map of their DEGs (Figure S4B) and another heat map of those expression values of top 30 amongst the most significant DEGs (Figure S4C). The impact of each of such DEGs (e.g., STC2, CBX2, ADAM1, and AKR1D1, Figure S4D) on the overall survival rate of HCC patients was evaluated by the Kaplan-Meier’s method and another single-gene-based Cox’s proportional hazards regression method (simply referred as to COX model). On this base, we have here established an eight-gene-based Cox’s prognostic model for HCC, with global p-value (< 0.01) and C-index (0.73) (Figure 1A), as well as the area under ROC curve (AUC=0.788, Figure 1B), manifesting a rather reliable performance of this model at predicting the overall survival rate (p=2.578e-08, Figure 1C). The survival status of patients grouped within high and low risk scores was further analyzed by the Kaplan-Meier’s method to construct their survival curves. The results revealed that the overall survival rate of the high-risk patients was significantly lower than that of those patients in the low-risk group (Figure 1C). By analyzing their survival time of two distinct risk groups, it was found that the number of death cases with the high risk scores was significantly higher than that of the low risk cases (Figure 1D). The expression levels of the model eight genes in the liver cancer tissues of distinct risk patients were illustrated in the heat map (Figure 1E), along with the specific parameters of this COX model being listed (in Table S4). Collectively, these demonstrate that the prognosis of HCC patients with distinct risks can be accurately predicted by this eight-gene-based prognostic model.

**Fig. 1.**
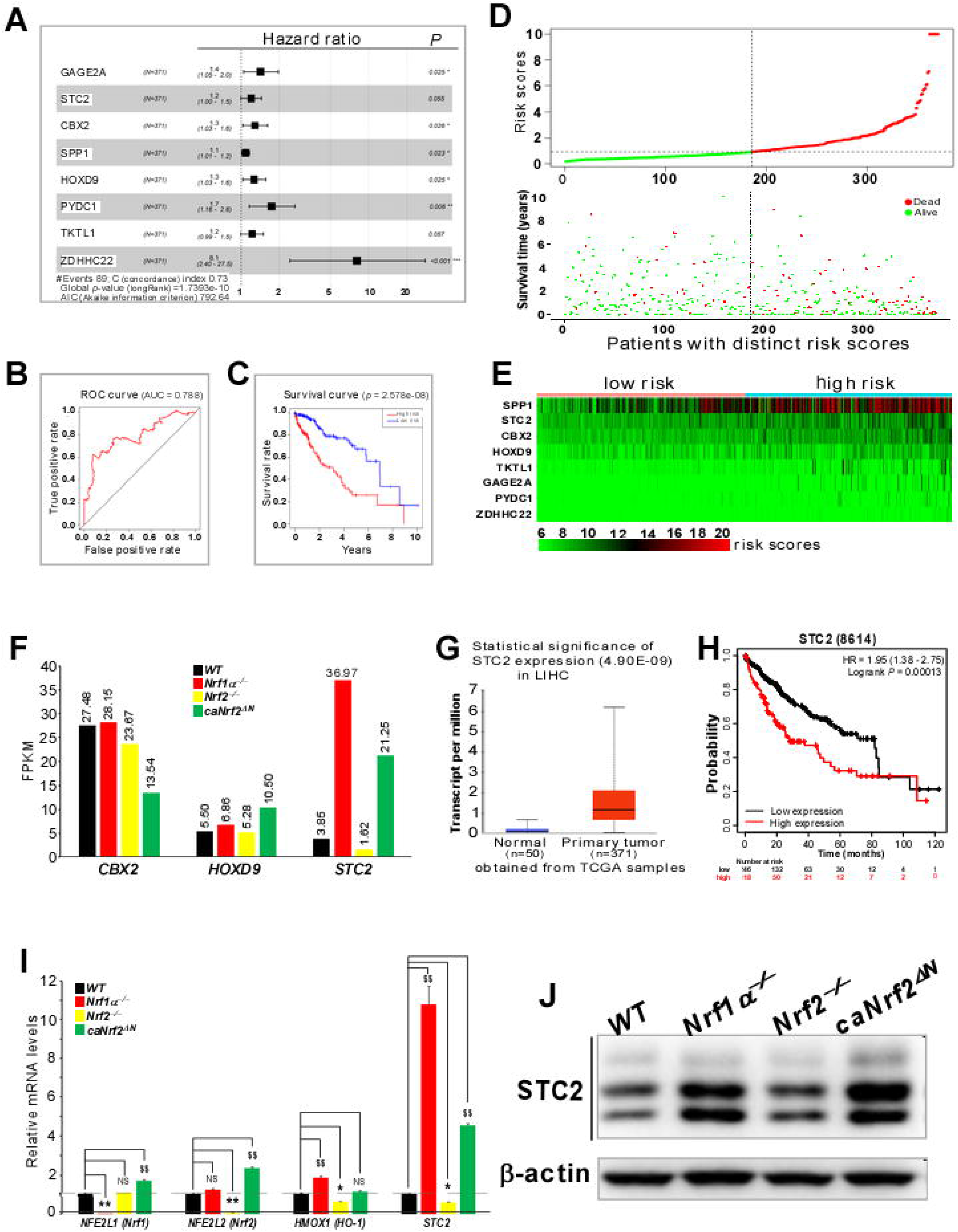
An involvement of STC2 in mediating the distinction between *Nrf1αNrf2α^-/-^* and Nrf2*^-/-^*. (A) A forest-map of the Hazard ratio of those genes included in the multi-gene prognostic model. (B) The ROC curve of the multi-gene prognostic model. (C) The overall survival rates of patients grouped within the high and low risks were analyzed by Kaplan-Meier. (D) Tow graphical representations of the COX model with the risk score of patients and their survival times. (E) A heat-map of the expression levels of eight genes indicated for the prognostic model of liver cancer tissues. (F) The mean FPKM values of CBX2, HOXD9 and STC2 expressed in *WT* (i.e., HepG2), and its derivative *Nrf1αNrf2α^-/-^*_Nrf2_*-/- Δ*N (G) The transcriptional expression levels of STC2 in liver cancer (LIHC) were obtained from the Ualcan database. (H) The effect of STC2 on the survival of HCC patients was analyzed by the Kaplan-Meier Plotter method. (I) The mRNA expression levels of STC2 in WT, *Nrf1αNrf2α^-/-^*, Nrf2*^-/-^* and caNrf2*^Δ^*^N^ cell lines were detected by RT-PCR. Data are reported as mean ± SD (n = 3 × 3, *p < 0.05, **p < 0.01, $$ p < 0.01, NS = no statistical difference). (J) The protein abundances of STC2 in WT, *Nrf1αNrf2α^-/-^*, Nrf2*^-/-^* and caNrf2*^Δ^*^N^ cell lines were determined by Western blotting with its specific antibody.

By comparative transcriptomic analysis of significant DEGs in the TCGA-LIHC tissues with those selected in Nrf1α, Nrf2 or caNrf2 (versus WT) cell lines (Figures S5 & S6), we found distinct expression levels of the COX-modeled genes (but only STC2 with a probability of 1.00) in each indicated cell line (Tables S5-S7). As a result, the predicted risk score of each cell line (Table 1) was calculated by multiplying all those gene coefficients by all their expression levels. Of note, a significant increase in the risk score was caused by loss of *Nrf1αNrf2α^-/-^* in HepG2 cells, but the risk score was markedly reduced by loss of Nrf2*^-/-^*; this seems to be consistent with discrepant phenotypes of their xenograft tumors in nude mice as described previously [15]. Further examination of the COX-modeled eight genes unraveled that GAGE2A, SPP1, TKTL1, ZDHHC22, PYDC1 were very less or not expressed in all four examined cell lines, while the expression levels of CBX2 and HOXD9 (albeit both may also serve markers) in *Nrf1αNrf2α^-/-^* or Nrf2*^-/-^* cell lines were not obviously different from those in *WT* cell line (Tables S5-S7, and Figure S6,E & F). However, it is, to our surprise, discovered that STC2 was significantly up-regulated in both cell lines of *Nrf1αNrf2α^-/-^* (retaining hyper-active Nrf2) and *^Δ^*^N^, but significantly down-regulated in Nrf2*^-/-^* cells as compared with that in *WT* cells (Figure 1F). Accordantly, STC2 was also significantly up-expressed in HCC when compared to the normal liver tissues by the Ualcan database [47]. Such increased expression of STC2 appears to presage a striking reduction in the overall survival rate of HCC patients (Figure 1, G & H), by using the Kaplan-Meier Plotter database [48]. Next, the mRNA and protein abundances of STC2 in all examined WT, *-/- ^-/-^* and *^Δ^*^N^ cell lines were further validated by RT-qPCR and western blotting, respectively. As expected, the results demonstrated that both mRNA and protein expression levels of STC2 were significantly up-regulated in *Nrf1αNrf2α^-/-^* and caNrf2 cell lines, whereas obviously down-expressed STC2 mRNA levels were determined in Nrf2*^-/-^* cells, but with an exception of no obvious changes in its protein expression, as compared with their control values obtained from *WT* cells (Figure 1, I & J). Herein, it is noteworthy that the validity of STC2 antibody was also verified by thapsigargin (TG)-stimulated expression of endogenous STC2 proteins, which was manifested with three distinct isoforms between 34-kDa and 39-kDa (Figure S1C). Altogether, it is inferable that STC2 is likely implicated in mediating the distinction between *Nrf1αNrf2α^-/-^* and *^-/-^*.

**Table 1.** Predicted risk scores of distinct hepatoma cell lines.

| Cell lines | Predicted risk scores |
| --- | --- |
| *WT* | 9.211808799 |
| *Nrf1*$\alpha^{-/-}$ | 15.95171908 |
| *Nrf2*$^{-/-}$ | 7.745601931 |
| *caNrf2*$\Delta^N$ | 10.2258014 |

### 3.2. HIF1A-dependent expression of STC2 was affected by Nrf1α or Nrf2 in distinct genotypic cell lines

A gene expression profiling interactive analysis (GEPIA [49]) of the liver cancer database was herein subjected to evaluating whether changes in the expression levels of STC2 are correlated with Nrf1, Nrf2, HIF1A and AHR (aryl hydrocarbon receptor), since they all are required for cytoprotection against oxidative stress [33, 50]. As shown in Figure S7 (A to D), the mRNA expression levels of Nrf1 and two known downstream genes *GCLM* and PSMB7 were significantly correlated (R > 0.2, p < 0.01), while the mRNA expression abundances of Nrf2 and typical downstream gene GCLM, as well as Nrf1, were positively correlated. Similarly, the mRNA expression levels of STC2 also appeared to be significantly correlated with Nrf1, Nrf2, HIF1A and AHR (Figure S7, E to H). The latter two transcription factors (albeit with relatively lower R values than those of the former two factors) had been reported to enable for directly binding the promoter region of the STC2 gene and hence considered as its direct upstream regulators [35, 51].

In accordance to the ChIP-Atlas database, HIF1A can directly bind to the promoter region of 5-kb before and after its transcription start site of STC2 in HepG2 cells (Figure 2A), but no similar binding data for AHR, Nrf1 or Nrf2 or AHR were found in this database. Rather, by further comparison of another CHIP-sequencing data for Nrf1 (from the Encode database) binding to the promoter regions of *GCLM* or STC2 in HepG2 cells (Figure S8, cf. A1 with A2), it is suggested that Nrf1 has a DNA-binding activity to STC2 as similar to binding its downstream GCLM. By contrast, Nrf2 possesses a significantly strong binding activity to *GCLM* rather than STC2 (Figure S8, cf. B1 with B2).

**Fig. 2.**
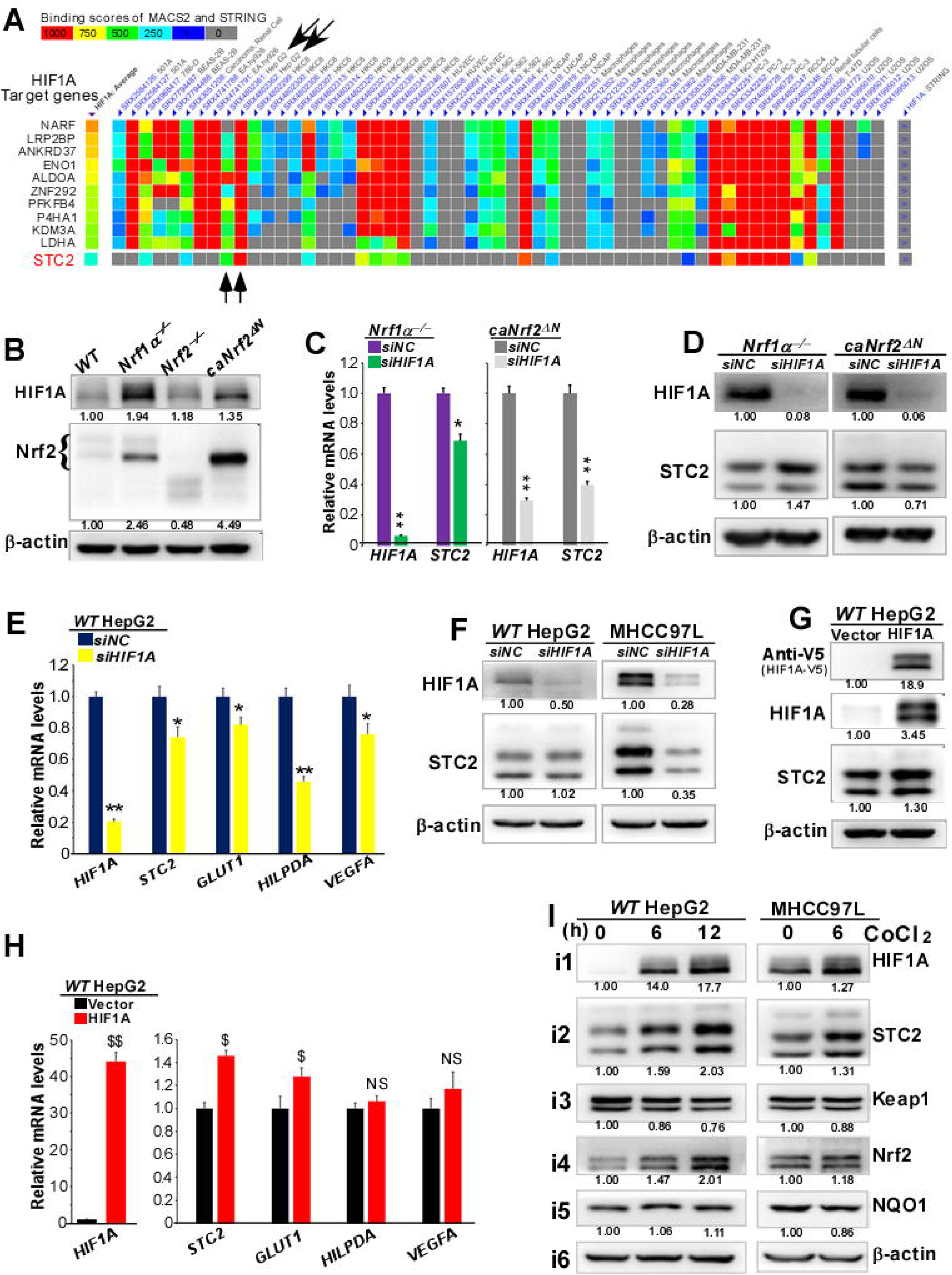
HIF1A-dependent expression of STC2 was affected by Nrf1α and/or Nrf2 in distinct genotypic cell lines. (A) HIF1A binds to the 5-Kbp promotor region adjoining the transcription start site of STC2 in HCC cells. The data were obtained from the ChIP-Atlas (at http://chip-atlas.org/). (B) The protein abundances of both HIF1A and Nrf2 in WT, *Nrf1αNrf2α^-/-^*, Nrf2*^-/-^* and caNrf2*^Δ^*^N^ cell lines were determined by Western blotting with their specific antibodies. (C) After transfection of HIF1A-targeting siRNA (siHIF1A) in *Nrf1αNrf2α^-/-^* and *^Δ^*^N^ cell lines, the mRNA levels of HIF1A and STC2 were examined by RT-qPCR. Data are reported as mean ± SD (n = 3 × 3, *p < 0.05, **p < 0.01). (D) After transfection of siHIF1A in *Nrf1αNrf2α^-/-^* and *^Δ^*^N^ cells, the protein abundances of HIF1A and STC2 were determined by Western blotting. (E) HepG2 cells were transfected with siHIF1A and then subjected to RT-qPCR analysis of the mRNA levels of HIF1A, STC2, GLUT1, HILPDA and VEGFA. Data are reported as mean ± SD (n = 3 × 3, *p < 0.05, **p < 0.01). (F) Both HepG2 and MHCC97L cell lines were transfected with siHIF1A, the protein levels of HIF1A and STC2 were detected by Western blotting. (G) HepG2 cells were transfected with a HIF1A-expressing construct and then subjected to immunoblotting analysis of the protein expression levels of both HIF1A and STC2. (H) The mRNA levels of HIF1A, STC2, GLUT1, HILPDA and VEGFA in HepG2 cells transfected with HIF1A-expressing construct were analyzed by RT-qPCR. Data are reported as mean ± SD (n = 3 × 3, $ p < 0.05, $$ p < 0.01, NS = no statistical difference). (I) Both HepG2 and MHCC97L cell lines were treated with (10μM dose of) cobalt chloride CoCl_2_ for 6 h or 12 h, and then subjected to Western blotting analysis of HIF1A, STC2, Keap1, Nrf2 and NQO1.

The expression abundances of HIF1A and Nrf2 were further validated by Western blotting of distinct genotypic cell lysates, as the resulting data revealed that both factors were significantly highly expressed in *Nrf1αNrf2α^-/-^* and caNrf2*^Δ^*^N^ cell lines (Figure 2B). Next, the real-time qPCR analysis unraveled that, upon silencing of HIF1A in *Nrf1αNrf2α^-/-^* or caNrf2*^Δ^*^N^ cell lines, its downstream STC2 expression levels were markedly down-regulated by siHIF1A in caNrf2*^Δ^*^N^ cells, but partially decreased by siHIF1A in *Nrf1αNrf2α^-/-^* cells (albeit with hyper-active Nrf2 accumulation) (Figure 2C). Similarly, the protein abundance of STC2 was significantly suppressed by siHIF1A in caNrf2*^Δ^*^N^ cells, but conversely elevated by knockdown of HIF1A in *Nrf1αNrf2α^-/-^* cells (Figure 2D). Such nuanced expression levels of STC2 imply a differential or even opposing response of this hormone to silencing of its upstream regulator HIF1A in distinct contexts between caNrf2*^Δ^*^N^ and *Nrf1αNrf2α^-/-^* cell lines.

Further examination of HIF1A-silenced *WT* HepG2 cells by real-time qPCR revealed that its downstream genes STC2, VEGFA (vascular endothelial growth factor A) and GLUT1 (glucose transporter 1, also called SLC2A1 (solute carrier family 2 member 1) were only marginally down-regulated by siHIF1A, except that HILPDA (hypoxia inducible lipid droplet associated) was significantly repressed by silencing of HIF1A (Figure 2E). Such being the case, almost no changes in the protein expression of STC2 were observed in siHIF1A-treated HepG2 cells, but the abundance of STC2 in MHCC97L (from a low metastatic HCC) cell line was markedly down-regulated by silencing of HIF1A (Figure 2F). Collectively, such differential responses of STC2 (along with other downstream genes) to silencing of HIF1A in different types of cells demonstrate that it may also be regulated by another HIF1A-independent pathway.

Forced expression of HIF1A in *WT* HepG2 cells led to increased expression of STC2 at its protein and mRNA levels (Figure 2,G & H), while the mRNA levels of GLUT1 were modestly increased by overexpression of HIF1A, but with almost no changes in the mRNA abundances of VEGFA and HILPDA (Figure 2H). Next, to address such distinct responses of STC2 and other HIF1A-target genes, HepG2 and MHCC97L cell lines were treated with cobalt chloride (CoCl_2_, as a hypoxia inducer to stabilize endogenous HIF1A [52, 53]). As anticipated, the results revealed significant increases in the protein abundances of HIF1A and STC2 following CoCl_2_ treatment of HepG2 and MHCC97L cell lines for 6 h or 12 h (Figure 2I). However, it was, much to our surprise, found that such CoCl_2_-stimulated expression of HIF1A and STC2 was also accompanied by significant increases of Nrf2, but with obvious decreases of its negative regulator Keap1 (Figure 2I, cf. i3 with i4). Taken altogether, these indicate that the transcriptional expression of STC2 and/or HIF1A may also be regulated by Nrf2, except that all these protein expression levels are, de facto, tightly controlled by Nrf1α-target proteasomes.

### 3.3. Distinct roles of Nrf2 and Nrf1α for regulating the expression of STC2 in distinct genotypic contexts

The above-described data suggested a HIF1-independent mechanism accounting for differential up-regulation of STC2 between *Nrf1αNrf2α^-/-^* and *^Δ^*^N^ cell lines may also exist. To gain an insight into this, we first compared the transcriptomic data of two HEK293 cell lines, that had been allowed for the tetracycline-inducibly stable expression of Nrf1α or Nrf2, respectively [19]. As shown in Figure 3A, a significant increase in the expression of STC2, but not of STC1, was determined in Nrf2-, rather than Nrf1α-, expressing cell lines, even although their co-target HO-1 was up-regulated in both cell lines, when compared with that of *WT* cells. Next, inducible increases in the endogenous expression of Nrf2, as well as its targets HO-1 and NQO1, in HepG2 cells MHCC97L cell lines were stimulated by oltipraz (as a known activator of Nrf2) (Figure 3B, b1, b4 & b5). Interestingly, such oltipraz-stimulated increase of Nrf2 was accompanied by a significant increment of STC2, along with another significant decrease of HIF1A, (Figure 3B, cf. b1, b2 with b3), all of which occurred at 24 h to 48 h after oltipraz treatment. Collectively, these demonstrate that except from HIF1A, Nrf2 is also required for mediating the transcriptional expression of STC2.

**Fig. 3.**
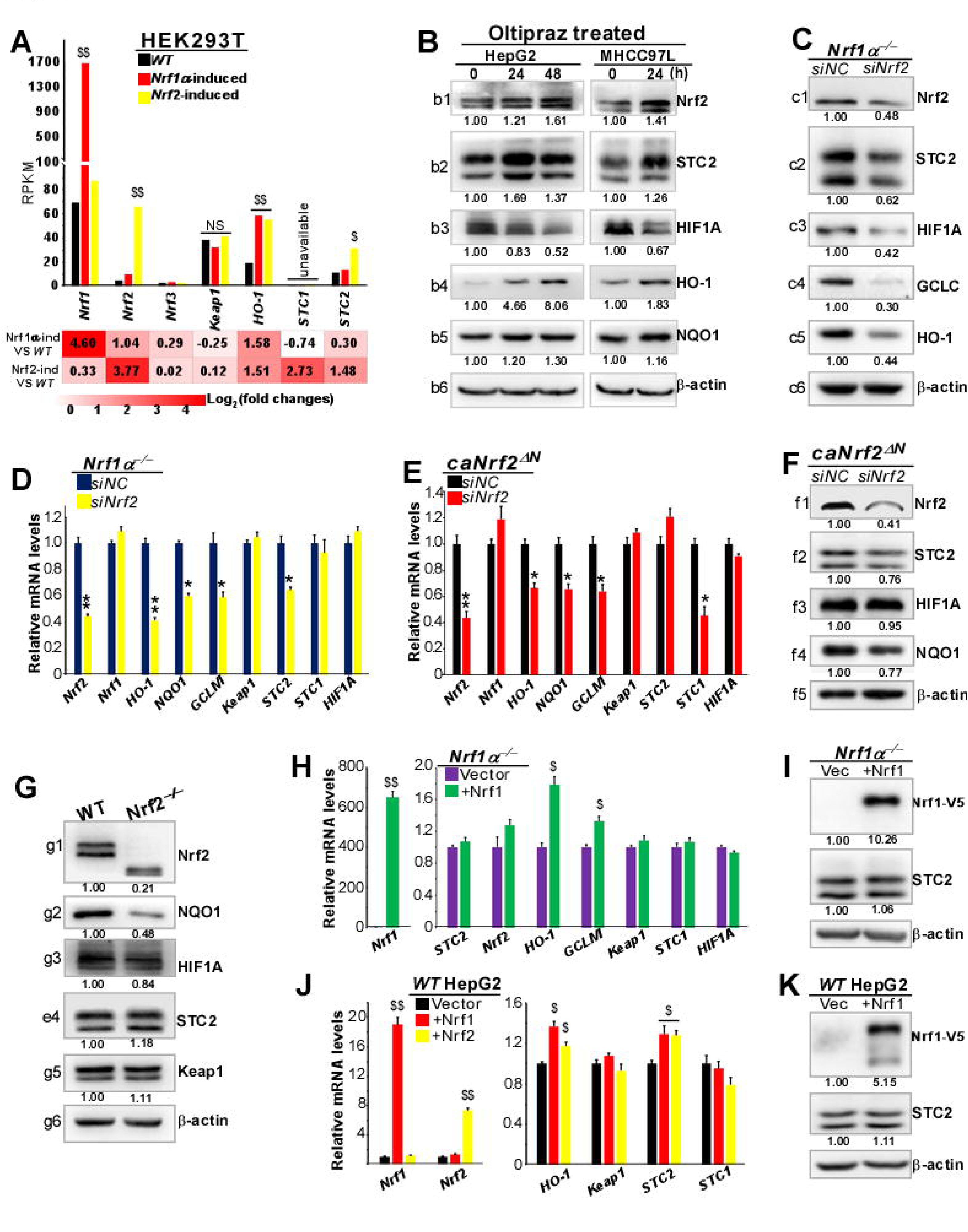
Distinct roles of Nrf2 and Nrf1α for regulating the STC2 expression in distinct genotypic contexts. (A) The expression RPKM values of STC2 and other indicated genes in Nrf1α- or Nrf2-induced HEK 293T cell lines were shown graphically (n = 3). (B) Both HepG2 and MHCC97L cell lines were treated with (10μM dose of) Oltipraz for 24 h or 48 h, and then subjected to Western blotting analysis of Nrf2, STC2, HIF1A, HO-1 and NQO1 proteins. (C) The Effect of Nrf2-targeting siRNA (siNrf2) on the protein expression levels of Nrf2, STC2, HIF1A, GCLC and HO-1 in *Nrf1αNrf2α^-/-^* cells was analyzed by Western blotting. (D) *Nrf1αNrf2α^-/-^* cells were transfected with siNrf2 and then subjected to RT-qPCR analysis of the mRNA levels of Nrf2, Nrf1, HO-1, NQO1, GCLM, Keap1, STC2, STC1 and HIF1A. Data are reported as mean ± SD (n = 3 × 3, *p < 0.05, **p < 0.01). (E) After transfection of caNrf2*^Δ^*^N^ cells with siNrf2, the mRNA levels of Nrf2, Nrf1, HO-1, NQO1, GCLM, Keap1, STC2, STC1 and HIF1A were determined by RT-qPCR. Data are reported as mean ± SD (n = 3 × 3, *p < 0.05, **p < 0.01). (F) The effect of siNrf2 on the protein expression of Nrf2, STC2, HIF1A and NQO1 in caNrf2*^Δ^*^N^ cells was detected by Western blotting. (G) The protein abundances of Nrf2, NQO1, HIF1A, STC2 and Keap1 in *WT* and Nrf2 *^-/-^* cell lines were detected by Western blotting (H) *Nrf1αNrf2α^-/-^* cells were transfected with a Nrf1-expressing plasmid, and then subjected to RT-qPCR detection of the mRNA levels of Nrf1, STC2, Nrf2, HO-1, GCLM, Keap1, STC1 and HIF1A. Data are reported as mean ± SD (n = 3 × 3, $ p < 0.05, $$ p < 0.01). (I) After overexpression of Nrf1 was allowed in *Nrf1αNrf2α^-/-^* cells, subsequent changes of both STC2 and Nrf1 proteins were determined by Western blotting. (J) HepG2 cells were transfected with Nrf1 or Nrf2 expression constructs and then subjected to RT-PCR analysis of the mRNA levels of Nrf1, Nrf2, HO-1, Keap1, STC2 and STC1, as shown as mean ± SD (n = 3 × 3, $ p < 0.05, $$ p < 0.01). (K) After Nrf1 expression plasmid were transfected into HepG2 cells, the changes of both Nrf1 and STC2 protein abundances were examined by Western blotting.

Such putative role of Nrf2 in the regulation of STC2 was further corroborated by silencing of this CNC-bZIP factor in the subsequent experiments. As shown in Figure 3(C & D), knockdown of Nrf2 led to obvious decreases in the protein and mRNA expression levels of STC2 in *^-/-^* cells, along with decreased expression of those known Nrf2-target HO-1, GCLC, *GCLM* and NQO1. Such being the case, HIF1A was also significantly reduced by siNrf2 at its protein abundance, but not its mRNA levels. By contrast, in caNrf2*^Δ^*^N^ cells the mRNA expression of STC2 was not decreased, but conversely modestly increased by siNrf2, while its protein abundance was partially down-regulated by silencing Nrf2 (Figure 3,E & F). However, both the mRNA and protein expression levels of HIF1A appeared to be unaffected by knockdown of Nrf2 in caNrf2*^Δ^*^N^ cells (but with a marginal increase of Nrf1 retained). Altogether, with the CHIP-sequencing data (from the Encode database) for binding of Nrf1 or Nrf2 to the promoter regions of STC2 and HIF1A (Figures S8 and S9A), these indicate distinct roles of Nrf2 and Nrf1 for monitoring the expression of STC2 and its upstream regulator HIF1A at distinct layers (from mRNA to protein levels) in different contexts.

When compared with *WT* cells, almost no changes in the basal expression of HIF1A and STC2 was observed in Nrf2*^-/-^* cells (Figure 3G, also see Figure 1J & 2B), implying another possible role of Nrf1 in regulating STC2. Yet, it is disappointing that the expression levels of STC2 and HIF1 were almost unaltered, although its target genes HO-1 and *GCLM* were up-regulated, by restoration of Nrf1 in *Nrf1αNrf2α^-/-^* cells (with an aberrant increase of Nrf2) (Figure 3,H & I). However, overexpression of Nrf1 (and Nrf2) in *WT* HepG2 cells resulted in marked increases in the mRNA expression of STC2 as well as HO-1 (Figure 3J), but with no obvious changes in the STC2 protein expression (Figure 3K). Lastly, a series of luciferase reporter assays unraveled that Nrf1, Nrf2 and HIF1A enabled distinct lengths of STC2 promoter-driven genes to be trans-activated (Figure S9B). Collectively, these demonstrate that like Nrf2, Nrf1 is involved in mediating the transcriptional expression of STC2, but in the meantime, its basal protein expression abundance may be also further monitored, to a certain constant extent, by Nrf1-target proteasomes in a negative feedback regulatory loop.

### 3.4. STC2 mediates a feedback regulatory loop to monitor the expression of HIF1A and Nrf2

Since the aforementioned data have manifested that Nrf1 and Nrf2 enable to promote differential expression levels of STC2 in HIF1-dependent and -independent fashions, thus we investigated whether there exists a feedback regulatory mechanism accounting for STC2 to maintain the redox homeostasis system involving HIF1A, Nrf1, Nrf2 and Keap1. As shown in Figure 4A, a significant increase of Keap1 resulted from silencing the expression of STC2 by siSTC2 in *WT* HepG2 cells; this was accompanied by marked decreases of Nrf2 and its downstream HO-1 (cf. a1 to a4). Interestingly, such silencing of STC2 also led to a striking decrease of HIF1A, but not Nrf1 (Figure 4B). Further real-time qPCR analysis revealed that the mRNA expression of HIF1A was significantly suppressed by siSTC2, along with partial down-regulation of HO-1 and *GCLM* (Figure 4C). However, it is intriguing that almost no changes in the mRNA levels of Nrf2 and Keap1, but with a significant increment of Nrf1. Collectively, these demonstrate a positive feedback loop between STC2 and HIF1. But those disparities between the protein and mRNA expression levels of Nrf1, Nrf2, Keap1 and their co-target HO-1 in the STC2-silenced HepG2 cells suggest at least two distinct feedback regulatory mechanisms at distinct strata (from mRNA to protein) existing among them within a multi-hierarchical endogenous network.

**Fig. 4.**
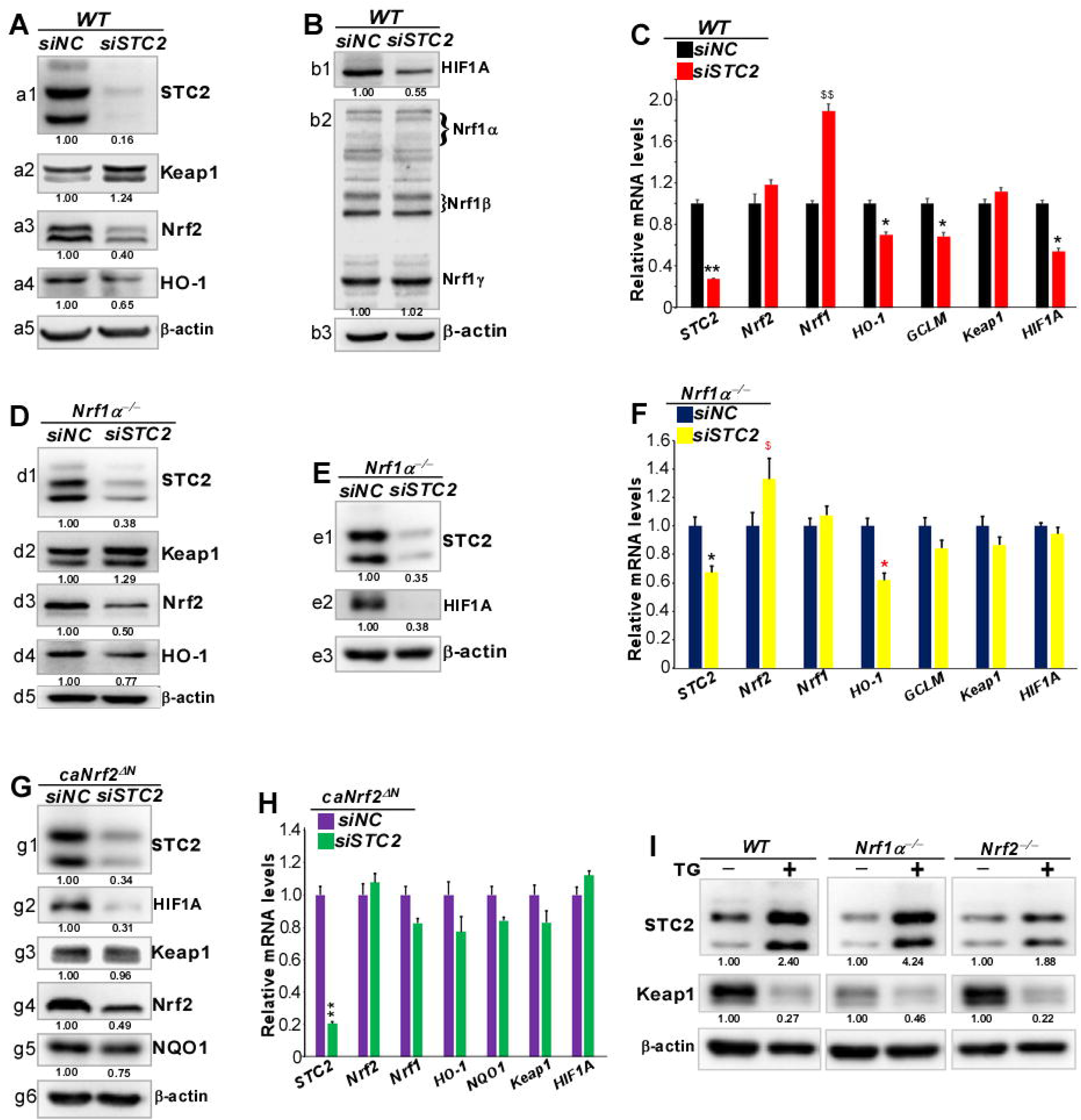
STC2 monitors the expression of HIF1A and Nrf2 through a putative feedback regulatory loop. (A) HepG2 cells were transfected with STC2-targeting siRNA (siSTC2), and then subjected to Western blotting analysis of STC2, Keap1, Nrf2 and HO-1 protein levels. (B) The protein levels of HIF1A and Nrf1 in siSTC2-transfected HepG2 cells were detected by Western blotting. (C) The mRNA expression levels of STC2, Nrf2, Nrf1, HO-1, GCLM, Keap1 and HIF1A in siSTC2-transfected HepG2 cells, were determined by RT-qPCR, and shown as mean ± SD (n = 3 × 3, *p < 0.05, **p < 0.01, $$ p < 0.01). (D) Western blotting analysis of STC2, Keap1, Nrf2 and HO-1 protein levels in siSTC2-transfected *Nrf1αNrf2α^-/-^* cells. (E) The effect of siSTC2 on HIF1A (and STC2) protein abundance in siSTC2-transfected *^-/-^* cells was determined by Western blotting. (F) The mRNA levels of STC2, Nrf2, Nrf1, HO-1, *GCLM*, Keap1 and HIF1A in siSTC2-transfected *Nrf1αNrf2α^-/-^* cells*-/-*Nrf1α cells were detected by RT-qPCR. Data are reported as mean ± SD (n = 3 × 3, *p < 0.05, $ p < 0.05). (G) Distinct effects of siSTC2 on STC2, HIF1A,Keap1, Nrf2 and NQO1 proteins in caNrf2*^Δ^*^N^ cells were examined by Western blotting. (H) The mRNA levels of STC2, Nrf2, Nrf1, HO-1, NQO1, Keap1 and HIF1A in siSTC2-transfected *^Δ^*^N^ cells were determined by RT-qPCR and shown graphically as mean ± SD (n = 3 × 3, **p < 0.01). (I) The inhibitory effect of Thapsigargin (TG) on the protein expression of Keap1 and STC2 was detected by Western blotting after HepG2 cells had been treated with this chemical (1μM dose).

Further examination of STC2-silenced *^-/-^* cells also unraveled that the protein abundances of Nrf2 and HO-1 were significantly decreased by siSTC2, as accompanied by a significant increase of Keap1 (Figure 4D, cf. d1 to d4). This occurred concomitantly with almost complete abolishment of HIF1 by siSTC2 (Figure 4E, cf. e2 with e1), but its mRNA expression appeared to be unaffected by siSTC2 (Figure 4F). Moreover, it is also hard to understand that siSTC2 led to a modest increase in the mRNA levels of Nrf2, but with a modest decrease of HO-1 Nrf1α cells (with hyper-active Nrf2 retained), while the mRNA levels of Keap1, along with the remnant Nrf1, were roughly unaltered by siSTC2 (Figure 4F). These ostensibly contradictory results suggest that the STC2 signaling feedback to HIF1A, Nrf1, Nrf2 and Keap1 is much likely to occur predominantly at their protein rather than mRNA strata.

By contrast, silencing of the STC2 expression in caNrf2*^Δ^*^N^ cells led to significant decreases in the protein levels of HIF1A, Nrf2 and NQO1, but with no obvious changes in the abundance of Keap1 (Figure 4G). In addition, their mRNA expression levels were largely unaffected by siSTC2 (Figure 4H). Strikingly, induction of the endogenous STC2 expression by TG (a microsomal Ca^2+^-ATPase inhibitor) in all examined WT,*-/-* _and_*-/-* cell lines resulted in significant decreases of Keap1 (Figure 4I). Altogether, these results demonstrate that STC2 mediates a feedback regulatory loop to promote the protein expression of Nrf2, as well as HIF1A, by antagonizing Keap1 and/or via a putative Ca^2+^-mediated signaling pathway. This is due to a fact that TG, as a classic endoplasmic reticulum stressor, can inhibit the transport of free Ca^2+^ and hence increase the intracellular Ca^2+^ levels [54], so that the Ca^2+^-mediated signaling was activated and/or prolonged insomuch as to monitor STC2 and Keap1.*<ι><ιτ></i>*

### 3.5. STC2 augments hepatoma cell proliferation and its malgrowth in vitro and in vivo

To clarify the biological role of STC2 in HCC, its gene-editing by CRISPR/Cas9 in HepG2 cells was employed to establish two mutant cell lines, designated STC2^insC^ and *STC2^-/-^*, that were further confirmed by their genomic DNA-sequencing (Figure S2A), real-time qPCR and Western blotting (Figure 5,A & B). By contrast with *STC2^-/-^*, STC2 remained to yield the smallest polypeptide of STC2 among its three distinct isoforms (Figure 5B), which is similar to the minor polypeptide arising from a mutant of STC2 (at the first translation starting codon into CTG, Figure S2C, #1). Another stably STC2-expressing cell line was established using a lentiviral system, and the expression efficiency of STC2 were further identified by Western blotting (Figure 5C) and real-time qPCR (Figure 5D). Of note, one cell line with its better validated effects was selected and named Lentiv-STC2 to use for the subsequent experiments.

**Fig. 5.**
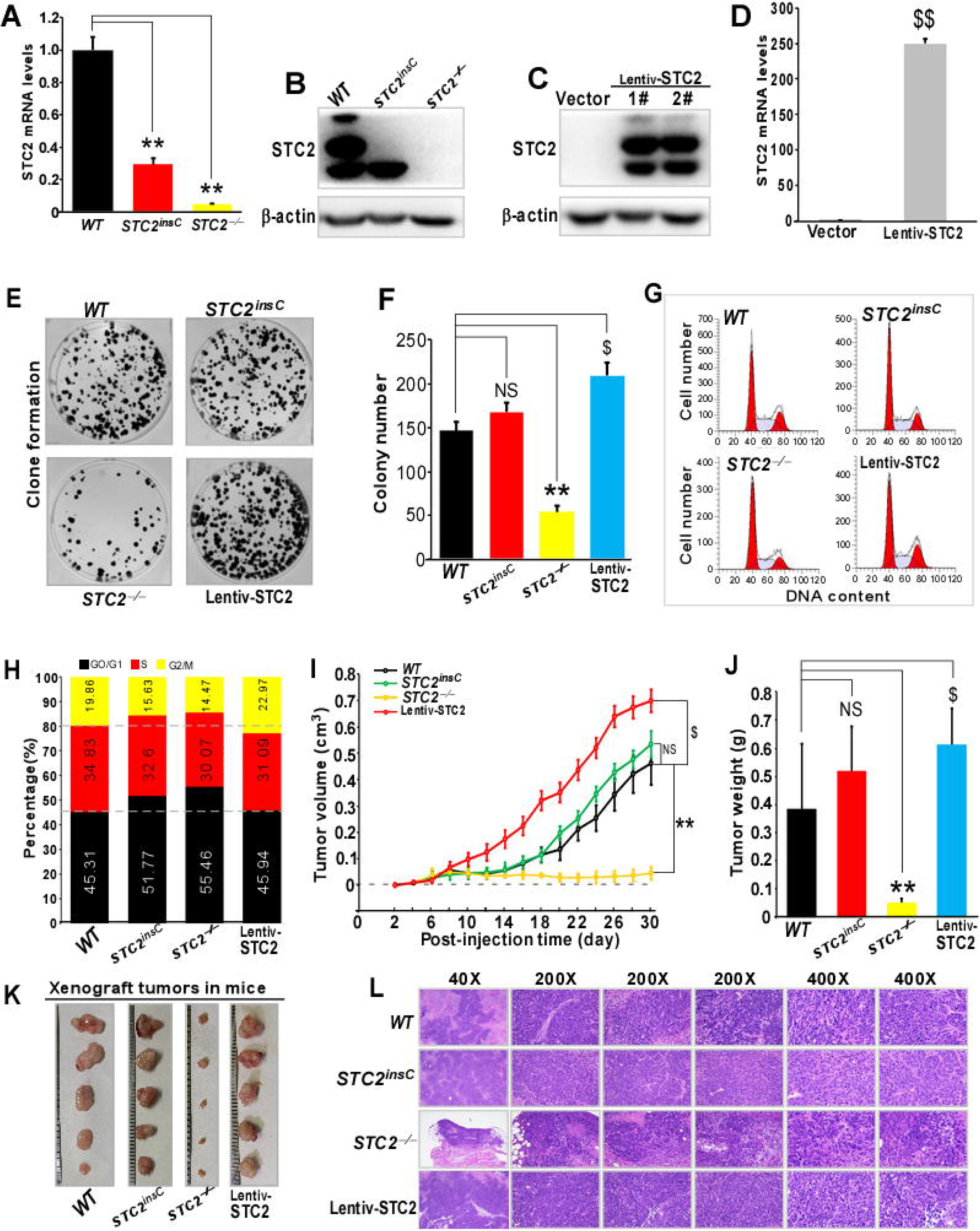
STC2 augments hepatoma cell proliferation and its malgrowth *in vitro* and *in vivo*. (A) The mRNA levels of STC2 in WT, STC2^insC^ and *STC2^-/-^* cell lines were determined by RT-qPCR and shown as mean ± SD (n = 3 × 3, **p < 0.01). (B) The protein levels of STC2 in WT, STC2^insC^ and *STC2^-/-^* cell lines were examined by Western blotting. (C) Western blotting analysis of the STC2 protein in HepG2 cells that had been transfected with the Lentiv-STC2 (#1 and #2) or an empty vector. (D) RT-qPCR analysis of the STC2 mRNA levels in HepG2 cells that had been transfected with Lentiv-STC2 (#1 and #2) or an empty vector. Data are reported as mean ± SD (n = 3 × 3, $$ p < 0.01). (E, F) Colony formation of WT, STC2^insC^, *^-/-^* and Lentiv-STC2 cell lines and their clone clusters were counted. Data are presented as mean ± SD (n =3; **p < 0.01, $ p < 0.05; NS = no statistical difference). (G,H) Distinct cell cycles were measured by flow cytometry. The data are obtained from two different experiments (n = 3) and shown graphically. (I) Different growth curves of mouse subcutaneous xenograft tumors derived from WT, STC2^insC^, *^-/-^* and Lentiv-STC2 cell lines and measured in size every two days, before being sacrificed on the 30^th^ day. Data are shown as mean ± SD (n = 5 per group, **p < 0.01; $ p < 0.05, NS = no statistical difference). (J) All those final tumor weights of distinct cell groups were calculated and shown as mean ± SD (n = 5, **p < 0.01; $ p < 0.05, NS = no statistical difference). (K) Representation of distinct xenograft tumors derived from WT, STC2^insC^, *^-/-^* and Lentiv-STC2 cell lines. (L) The histological photographs of indicated tumors were achieved by HE (hematoxylin & eosin) staining. Distinct scale bars = 500 µm in ×40 pictures, 100 µm in ×200 pictures and 50 µm in ×400 pictures.

Next, the biological functioning of STC2 in hepatoma cell growth and proliferation was assessed on the base of the above-established cell lines STC2^insC^, *STC2^-/-^* and Lentiv-STC2. As shown in Figure 5(E &F), the clone formation rate of HepG2 cells was almost completely suppressed by knockout of *STC2^-/-^*, but largely unaffected by the knockin mutant STC2 . By contrast, the colony formation rate of Lentiv-STC2 cells was significantly enhance by ectopically-expressing STC2 (Figure 5F). Subsequently, the changes in the cell-cycle of four distinct cell lines were determined by flow cytometry (Figure 5G). The results revealed that, when compared with the *WT* controls, *STC2^-/-^* and ^insC^ cell lines were significantly arrested at their G0/G1 phases, but conversely their

G2/M phases were thus shortened (Figure 5H), so as to enable the cell growth to be decelerated. By sharp contrast, the S-phase of Lentiv-STC2 cells was shortened, while its G2/M phase was accordingly lengthened (Figure 5H), such that the number of cells at the division phase increased, and the cell growth were thus accelerated (Figure 5,F & G). Collectively, these indicate that STC2 promotes the cell division and proliferation of hepatoma and its clonogenicity.

In order to further investigate the in vivo effect of STC2 on heptoma cell growth, the relevant xenograft models were established by subcutaneously injecting each of the indicated cell lines into nude mice. The cell proliferation in vivo was evaluated by measuring tumor volumes and weights. All the tumor sizes were measured every two days, until the 30^nd^ day when all mice were sacrificed and their transplanted tumors were then excised. As shown in Figure 5I, the results revealed that knockout of *^-/-^* resulted in a significant blockage of the tumor growth in mice, while Lentiv-STC2 overexpression enabled for promotion of its tumor malgrowth, but the STC2 -derived tumor growth changed negligibly, when compared with *WT* controls. Of note on the 30th day, the average volume and weight of those tumors derived from *STC2^-/-^* cells were substantially lowered than those from the *WT* controls (Figure 5J). By contrast, Lentiv-STC2-derived tumors were significantly larger in size and also heavier in weight than all other tumors (Figure 5, J & K). Moreover, histological examination unraveled a mass of the coagulative necrosis in *STC2^-/-^*-derived tumors, but not in other tumors (Figure 5L). Taken together, these results demonstrate that STC2 has a potent tumorigenicity in HCC to promote the cell proliferation and its malgrowth in vivo and in vitro.

### 3.6. Functional annotation of DEGs in STC2^-/-^ or Lentiv-STC2 versus *WT* cells by transcriptome sequencing

To gain an insight into the pathobiological role of STC2 in distinct hepatoma phenotypes, *STC2^-/-^*, Lentiv-STC2 and *WT* cell lines were further subjected to transcriptome sequencing. Figure S10(A & B) showed a correlation heatmap of all examined samples and another boxplot of their expression-quantified distribution, respectively. The TPM values of STC2 were determined in the examined cell lines (Figure 6A); this indicates that the STC2 mRNA levels in each sample were fully consistent with the results from transcriptome sequencing. Subsequently, all those differentially expressed genes (DEGs) were defined by detecting their expression levels of |Log_2_[fold changes]| ≥ 2, calibrated p-value (Q-value) ≤ 0.05 and diverged probability of ≥ 0.8, relative to equivalents measured from control cells (Figure 6B). Of note, 204 DEGs were upregulated and 222 DEGs were downregulated in *STC2^-/-^* cells compared with *WT* cells, while 234 genes were upregulated and 226 genes were downregulated in Lentiv-STC2 cells (Figure S10,C & D). In contrast with *STC2^-/-^*, Lentiv-STC2 cells were manifested with 257 of upregulated genes and 251 of downregulated genes (Figures 6B and S10E). In Venn diagram (Figure 6C), 149 DEGs were detected identically in both *STC2^-/-^* and Lentiv-STC2 cell lines, of which 50 DEGs were further scrutinized in distinct combinations of every two groups.

**Fig. 6.**
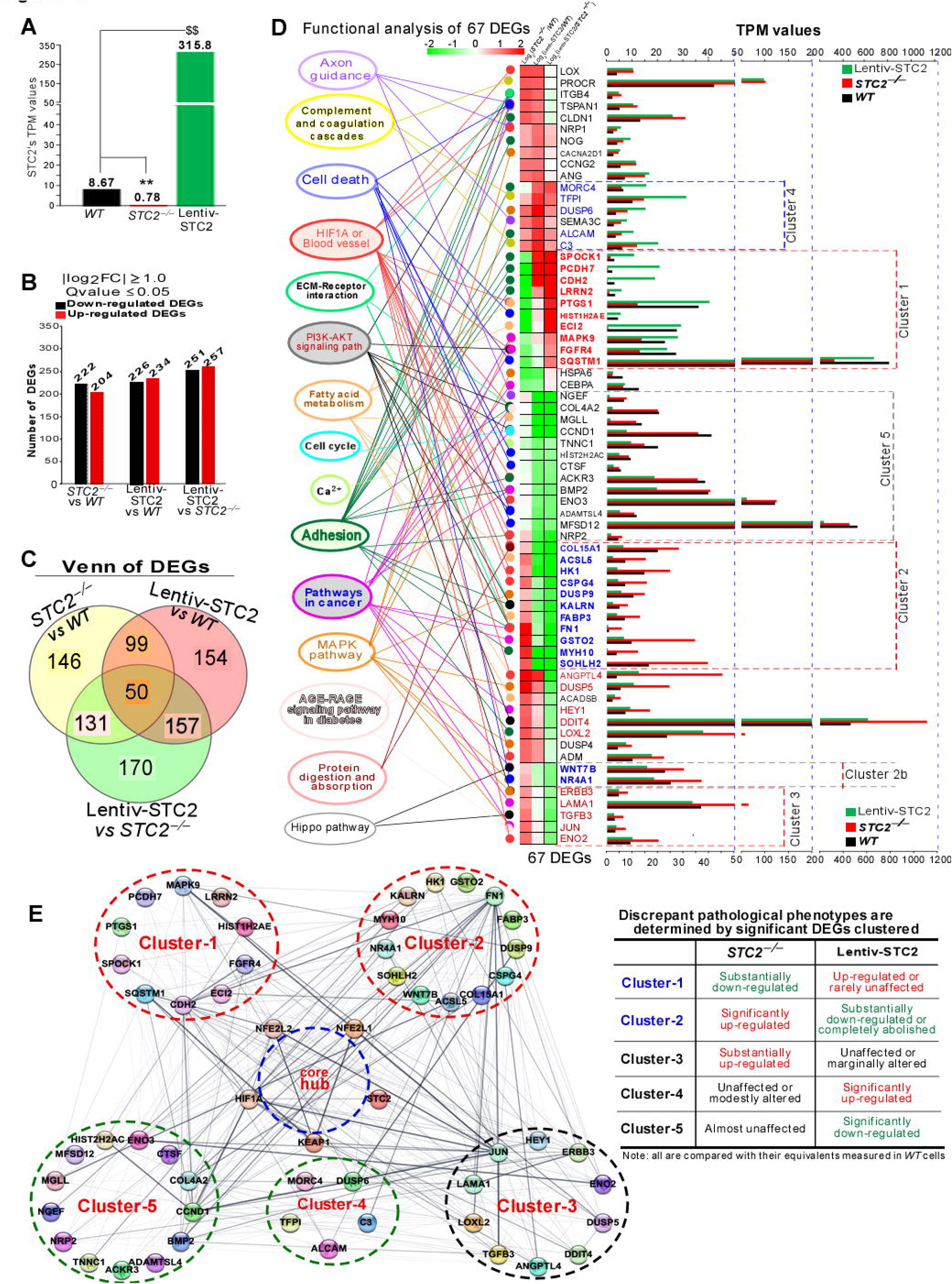
Transcriptome sequencing to identify DEGs significantly in *STC2^-/-^* or Lentiv-STC2 vs *WT* cell lines. (A) Graphical illustration of the STC2 expression at its TPM values (n = 3, **p < 0.01; $$, p < 0.01) in WT, *STC2^-/-^* and Lentiv-STC2 cell lines (B) Quantitative statistics of DEGs between every two groups of WT, *STC2^-/-^* and Lentiv-STC2 cell lines. (C) Venn diagram of DEGs between every two groups of WT, *^-/-^* and Lentiv-STC2 cell lines. (D) Significant DEGs of 67 (at their TPM values) with distinct functional annotation in important signaling pathways and key modules enriched in Lentiv-STC2, *STC2^-/-^* and *WT* cell lines. (E) Five major clusters of DEGs determine discrepant pathological phenotypes between *STC2^-/-^* and Lentiv-STC2 cells as compared with *WT* cells.

The terms of the biological process, cellular component and molecular function, along with putative pathways mediated by STC2, were annotated by enriching those significant DEGs based on both GO and KEGG and databases, respectively. The results were illustrated within histograms and scatterplots (Figure S11). The GO analysis revealed that the top 20 biological process terms of *STC2^-/-^* vs *WT* cells and Lentiv-STC2 vs *WT* cells corporately associated with angiogenesis, cell adhesion, multicellular organism development and extracellular matrix (ECM) organization. The cellular component terms associated with ECM, components of membrane, cytoskeleton and the endoplasmic reticulum lumen. The molecular function terms associated with structural constitution of ECM, oleamide hydrolase activity and anandamide amidohydrolase activity (Figure S11A). The KEGG enrichment analysis unraveled the main enrichments of those DEGs in axon guidance, protein digestion and absorption, advanced glycation endproducts (AGE)-RAGE (the receptor of AGE) signaling pathway in diabetics, fatty acid degradation, PI3K-AKT pathway, focal adhesion and ECM-receptor interaction (Figure S11B). These results indicate that STC2 has certain potential effects on substance-energy metabolism, extracellular signaling and cancer-related pathways. Overall, STC2 can influence relevant signaling and enzyme activity, as well as cell membrane structure, cytoskeleton and ECM, and thus affect cell proliferation and behavior, development and growth, and even pathological process.

By further scrutinizing those critical genes for those important pathways and functional modules significantly enriched for DEGs in *STC2^-/-^*, Lentiv-STC2 vs *WT* cell lines, such key genes of 67 were carefully selected and their expression levels with their functional enrichments of 15 were shown (in Figure 6D). Among them, the core Cluster-1 genes of 10 were substantially down-regulated in *STC2^-/-^*cells, but mostly up-regulated or rarely unaffected in Lentiv-STC2 cells, by comparison with those measured in *WT* cells. The Cluster-1 genes include MAPK9, HIST1H2AE (encoding histone H2AC8), CDH2 (cadherin 2), PCDH7 (protocadherin 7), LRRN2 (leucine rich repeat neuronal 2, a cell-adhesion molecule and/or signal transduction receptor), SPOCK1 (a proteoglycan to act as a protease inhibitor), PTGS1 (prostaglandin-endoperoxide synthase 1, also known as cyclooxygenase 1 [COX1]), ECI2 (enoyl-CoA delta isomerase 2), FGFR4 and SQSTM1 (i.e., p62, as a scaffolding protein required for autophagy of Keap1). Conversely, those Cluster-2 genes were significantly up-regulated in *STC2^-/-^*cells, but substantially down-regulated or even completely abolished in Lentiv-STC2 cells, when compared to their equivalents measured in *WT* cells. Such 13 of Cluster-2 genes include GSTO2 (glutathione S-transferase omega 2), SOHLH2 (encoding a bHLH transcription factor involved in spermatogenesis, oogenesis and folliculogenesis), COL15A1 (collagen 15α1 chain), CSPG4 (chondroitin sulfate proteoglycan 4, stabilizing a cell-substratum interaction on the endothelial basement membranes), KALRN (kalirin RhoGEF kinase, that interacts with HAP1 [huntingtin-associated protein 1] for vesicle trafficking), MYH10 (myosin heavy chain 10), FN1 (fibronectin 1), DUSP9 (dual specificity phosphatase 9, enabling for inactivation of its target MAPK family), HK1 (hexokinase 1), *ACSL5* (acyl-CoA synthetase long chain family member 5), and FABP3 (fatty acid binding protein 3, that participates in the long-chain fatty acid uptake, metabolism and transport), plus two Cluster-2b genes WNT7B (Wnt7B, a secreted signal to regulate cell fate and patterning in embryogenesis, oncogenesis and developmental processes) and NR4A1 (nuclear receptor 4A1, that serves as a transcription factor of the steroid-thyroid hormone-retinoid receptor superfamily to induce apoptosis after being translocated to the mitochondria).

Furtherly, those Cluster-3 genes were also substantially up-regulated in *STC2^-/-^*cells, but largely unaffected or marginally altered in Lentiv-STC2 cells, when compared with their equivalents of *WT* control cells (Figure 6D). Such 10 genes are ERBB3 (an EGFR family member called ErbB2-3 or HER3), LAMA1 (laminin subunit alpha 1, a portion of extracellular matrix glycoproteins), TGFB3 (transforming growth factor β3, a secreted ligand to bind various TGFβ receptors leading to recruitment and activation of SMAD family transcription factors that regulate gene expression), JUN (a proto-oncogene subunit of AP-1 transcription factor directly interacting with specific target genes), ENO2 (enolase 2, acting as an isoenzyme homodimer in mature neurons or cells of neuronal origin), DDIT4 (DNA damage inducible transcript 4, that negatively regulates the mTOR signaling in response to hypoxia, besides binding 14-3-3 protein), HEY1 (hair-like division-related enhancer 1, a bHLH transcription repressor of the HESR family required for embryonic development, neurogenesis and somitogenesis), DUSP5 (dual specificity phosphatase 5), LOXL2 (lysyl oxidase like 2, a member of the family essential for the biogenesis of connective tissue by catalyzing the first step in the formation of crosslinks in collagens and elastin), and ANGPTL4 (angiopoietin like 4, a secreted protein with a C-terminal fibrinogen domain to regulate glucose homeostasis, lipid metabolism, and insulin sensitivity).

Several genes were significantly up-regulated in Lentiv-STC2 cells, but roughly unaffected or modestly altered in *^-/-^*cells, when compared to their equivalent *WT* controls (Figure 6D). Such 5 genes (in Cluster-4) are MORC4 (a member of the MORC [microrchidia] family sharing an N-terminal ATPase-like ATP-binding region and a CW four-cysteine zinc-finger motif, also with a nuclear matrix binding domain and a two-stranded coiled-coil motif near its C-terminus), TFPI (tissue factor pathway inhibitor, serves a Kunitz-type serine protease inhibitor to regulate the tissue factor-dependent pathway of blood coagulation), DUSP6 (dual specificity phosphatase 6), ALCAM (activated leukocyte cell adhesion molecule, also known as CD166 [cluster of differentiation 166]), and C3 (complement C3, playing a central role in the activation of complement system). Conversely, 13 genes (in Cluster-5) were significantly down-regulated in Lentiv-STC2 cells, but almost unaffected STC2 cells, when compared to those equivalents in *WT* cells (Figure 6D). They were MFSD12 (major facilitator superfamily domain containing 12, that enables cysteine transmembrane transporter activity and regulates melanin biosynthesis and pigment metabolism), ENO3 (enolase 3, involved in muscle development and regeneration), BMP2 (bone morphogenetic protein 2, a secreted ligand of the TGF-β superfamily that binds its receptors leading to recruitment and activation of SMAD family transcription factors), ACKR3 (atypical chemokine receptor 3, a G-protein-coupled receptor family member), CCND1 (cyclin D1, as a regulatory subunit of CDK4 or CDK6), COL4A2 (collagen 4α2 chain), NGEF (neuronal guanine nucleotide exchange factor), MGLL (monoglyceride lipase), TNNC1 (troponin C1, a subunit of troponin exerting a central role in striated muscle contraction by binding calcium to abolish the inhibitory action, allowing actin interaction with myosin to generate tension), HIST2H2AC (histone H2AC), CTSF (cathepsin F, a cysteine proteinase of papain family serving as a major component of the lysosomal proteolytic system), ADAMTSL4 (the ADAMTS [a disintegrin and metalloproteinase with thrombospondin motifs]-like gene family member 4, with seven thrombospondin type 1 repeats that may exert diverse roles in cell adhesion, angiogenesis, and the developing nervous patterning), and NRP2 (neuropilin 2, a transmembrane protein that binds to SEMA3C and SEMA3F proteins and interacts with VEGF).

Collectively, these demonstrate that distinct pathological phenotypes of between *STC2^-/-^*- and Lentiv-STC2-derived hepatoma cell lines are determined principally by their key DEGs in Cluster-1 and Cluster-2 (Figure 6E). The *STC2^-/-^* defective phenotype was also strengthened by alterations of its specific genes in Cluster-3, while the Lentiv-STC2-expressing phenotype was further enhanced by changes in its specific gene expression profiling of Cluster-4 and Cluster-5 (Figure 6E, right panel). In addition to a Pearson correlation analysis of those core genes expressed in all the examined cell lines (Figures 7A and S12), the relativity between those cell lines was also evaluated (Table S8). As expected, the results unveiled that, on a whole, Lentiv-STC2 overexpressing cell line has a closer relevance to *Nrf1αNrf2α^-/-^* or caNrf2*^Δ^*^N^ cell lines (both with hyper-expressed STC2), whereas *^-/-^* cell line is only slightly relevant to *^-/-^* cell line (albeit with a striking diminishment of STC2), but largely not to caNrf2*^Δ^*^N^ cells (Figure 7B). Since such distinct pathological phenotypes are determined by altered programming of key gene transcription to mRNA translation into proteins, those DEGs governing critical transcription factors (Figures S13 & S14) and Ca^2+^ signaling molecules (Figure S15) regulated by Nrf1, Nrf2 and STC2 were further scrutinized, respectively.

**Fig. 7.**
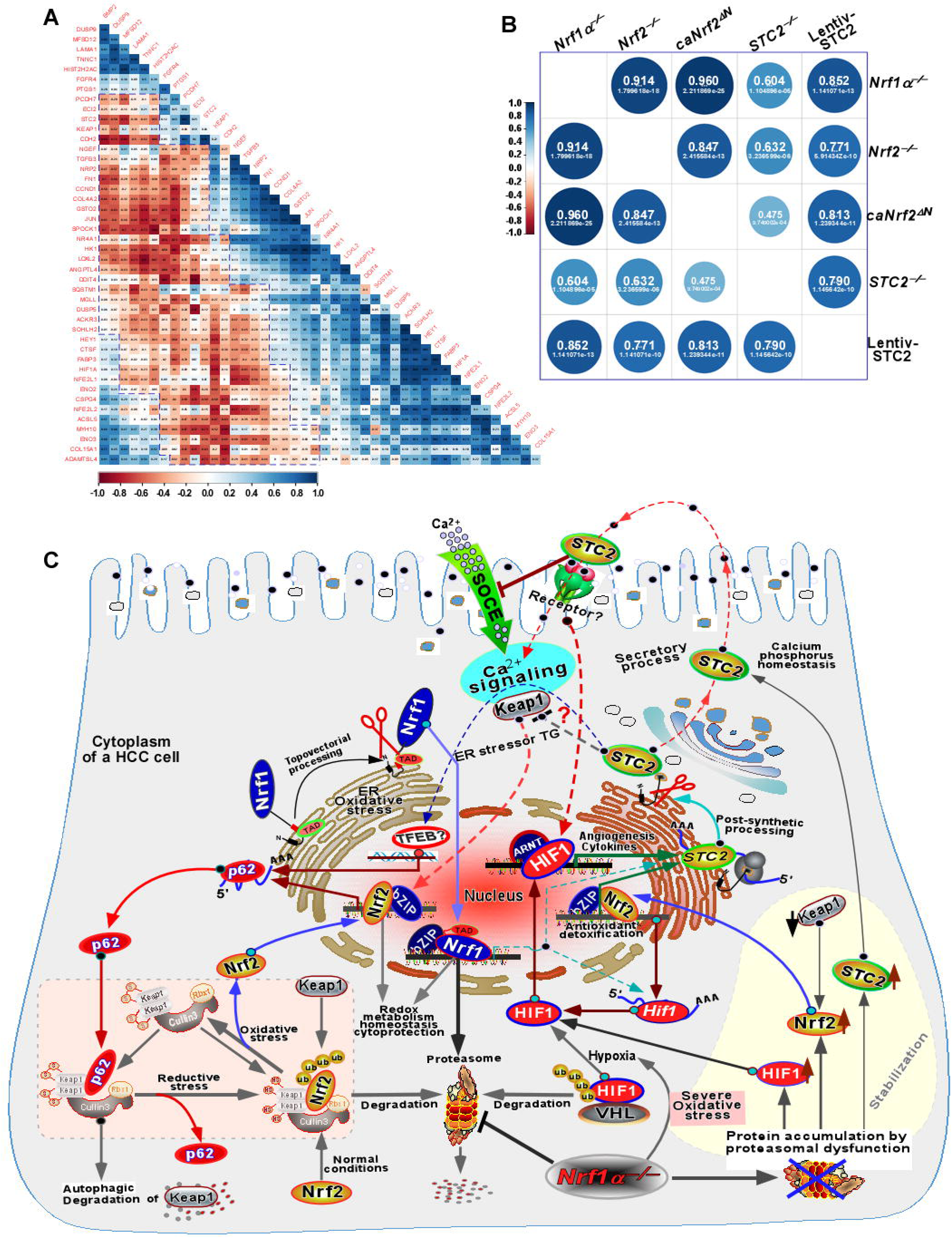
A model is proposed for a better understanding of STC2’s function in mediating discrepant phenotypes of between *Nrf1αNrf2α^-/-^* and Nrf2*^-/-^*. (A) The Pearson correlation analysis of key genes expressed significantly in WT, *-/--/-D*N *-/-* and Lentiv-STC2 cell lines, along with the relevant coefficients between every two of those genes. (B) The relevance coefficients between every two of*-/--/-^Δ^*^N^*^-/-^* and Lentiv-STC2 cell lines was calculated as shown (in big numbers) along with their p-values (in small numbers). (C) A model proposed for a better understanding of those key molecular inter-regulatory networks accounting for the role of STC2 in mediating distinctive phenotypes between *Nrf1αNrf2α^-/-^* and Nrf2*^-/-^*. Distinct intrinsic status of such a robust endogenous molecular-cellular network was further deciphered in Fig. 8.

## 4. Discussion

In the present study, we have established a prognostic model of liver cancer by mining the transcriptome data saved in the TCGA database and calculated the predictive risk scores of WT, *Nrf1αNrf2α^-/-^*, Nrf2*^-/-^* and caNrf2*^Δ^*^N^-derived tumors. The resulting evidence clearly demonstrates that loss of Nrf1α leads to a significant increase in the risk score, but the risk score is strikingly reduced by loss of Nrf2. Such distinction between Nrf1α and Nrf2 is fully consistent with our previously-reported phenotypic disparities of their xenograft tumors in nude mice [15]. Thus, based on the mathematic models of systems biology developed by Ao’s group [55-58], it is inferable that discrepant phenotypes between *Nrf1αNrf2α^-/-^* and Nrf2*^-/-^*-derived xenograft tumors are determined by different profiling of those key differential expression genes at distinct intrinsic status of a robust endogenous molecular-cellular network (as illustrated in Figure 7C). Therein, differential expression of a minimum set of key genes at distinct strata (e.g., from mRNAs to proteins) is de facto exhibited at their abundances, activities and topoforms in different phase transition, along with their intricate interactions between those core modular molecules in different subcellular contexts.

Of note, the expression of STC2 was upregulated in liver cancer tissues and also coincided with a reduction of the overall survival rate of patients with hepatomas. This is also completely consistent with those previous reports of STC2 being upregulated in multiple types of cancers [28, 59-61]. So highly up-regulated expression levels of STC2 in liver cancer are also associated with the poor prognosis of relevant patients [59-63]. Importantly, significant up-regulation of STC2 was examined in *Nrf1αNrf2α^-/-^* cells (with an aberrant Nrf2 accumulation) and caNrf2*^Δ^*^N^ cells (in which Nrf2 is constitutively activated owing to a loss of its N-terminal keap1-binding Neh2 domain). By contrast, a lower mRNA expression level of STC2 in *WT* cells was determined to be only about one-tenth of that measured in *Nrf1αNrf2α^-/-^* cells. However, down-regulated mRNA expression levels of STC2 in Nrf2*^-/-^* cells were accompanied by no significant changes in its protein levels when compared to *WT* controls, implying there exists a nonlinear stochastic feedback regulation of between mRNA and protein expression of STC2 by, at least, Nrf2 and its target genes. As such, these collective results demonstrate that STC2 is, as a potent biomarker for hepatocellular carcinoma, also implicated in mediating distinct phenotypes of between *Nrf1αNrf2α^-/-^* and Nrf2*^-/-^*-derived tumors.

STC2 has been widely accepted as a regulator of both calcium and phosphorus homeostasis, of which calcium ion (Ca^2+^, as the second messenger to initiate signaling networks) can regulate a variety of cellular processes, such as gene transcription, mRNA translation into protein, protein folding and quality control, cell metabolism, division and proliferation [64]. Interestingly, the inducible expression of STC2 is also evidently stimulated by oxidative stress and hypoxia [33], leading to a limitation of the STIM1-mediated store-operated Ca^2+^ entry (SOCE) into the triggered cells during cellular stress in order to promote cellular survival [44]. Our experimental evidence has been presented, together with another previous study [35], revealing that HIF1A is an upstream regulator of STC2 by directly binding the promoter of STC2, and this molecular event is also monitored by Nrf2 (Figures 7C & 8A), albeit HIF1A and Nrf2 are two known master regulators of hypoxia and oxidative stress, respectively [65, 66]. Besides, induction of STC2 is significantly stimulated by the endoplasmic reticulum stressor TG (as a microsomal Ca^2+^-ATPase inhibitor to cause an accumulation of Ca^2+^ in the oxidative lumen of this organelle required for the local mRNA translation into protein, and its quality control). The induction is inferable to be also accompanied by TG-stimulated expression of the redox-determining factor Nrf1 integrated in the endoplasmic reticulum [67]. Specific knockout of Nrf1α leads to an severe increase in the intracellular reactive oxygen species (ROS) [15, 68]. Such overproduction of ROS causes inactivation of PHD2 by oxidation of the ferrous ion essential for the central catalytic hydroxylation of prolines, so to inhibit the hydroxylation of HIF1α and hence stabilize its protein expression [69]. This, as a result, leads to the increased STC2 protein expression in *Nrf1αNrf2α^-/-^* cells. The stabilization of HIF1α protein is also reinforced by proteasomal dysfunction in *Nrf1αNrf2α^-/-^* cells (Figures 7C and 8B).

Intriguingly, we also found that the STC2 expression is promoted by Nrf2, independently of HIF1A. This is due to the supportive evidence showing that the expression of STC2 protein was significantly up-regulated by oltipraz, albeit the protein abundance of HIF1A was markedly inhibited by this inducer (of Nrf2 that had been shown to facilitate the ubiquitin-mediated degradation of HIF-1α [70, 71]), as accompanied by promoted expression levels of Nrf2 and its targets HO-1 and NQO1. Similarly, the abundance of STC2 was not reduced by silencing of HIF1A in *Nrf1αNrf2α^-/-^* cells (retaining hyper-active Nrf2 and HIF1A), albeit it was significantly down-regulated by knockdown of HIF1A by siHIF1A in caNrf2*^Δ^*^N^ cells. Besides, such a genomic loss of the N-terminal keap1-binding Neh2 domain in caNrf2*^Δ^*^N^ cells also enables prevention of putative Keap1-mediated degradation of this mutant factor, leading to the increased expression of STC2. However, a role for Nrf1 in augmenting STC2 and HIF1A cannot also be ruled out, because this CNC-bZIP factor is up-regulated in *^Δ^*^N^ cells (Figure 8C). Further examinations revealed that Nrf1 and Nrf2 can bind to the promoter region of STC2, as well as HIF1A, and also mediate its transcriptional expression (Figures S8 and S9).

**Fig. 8.**
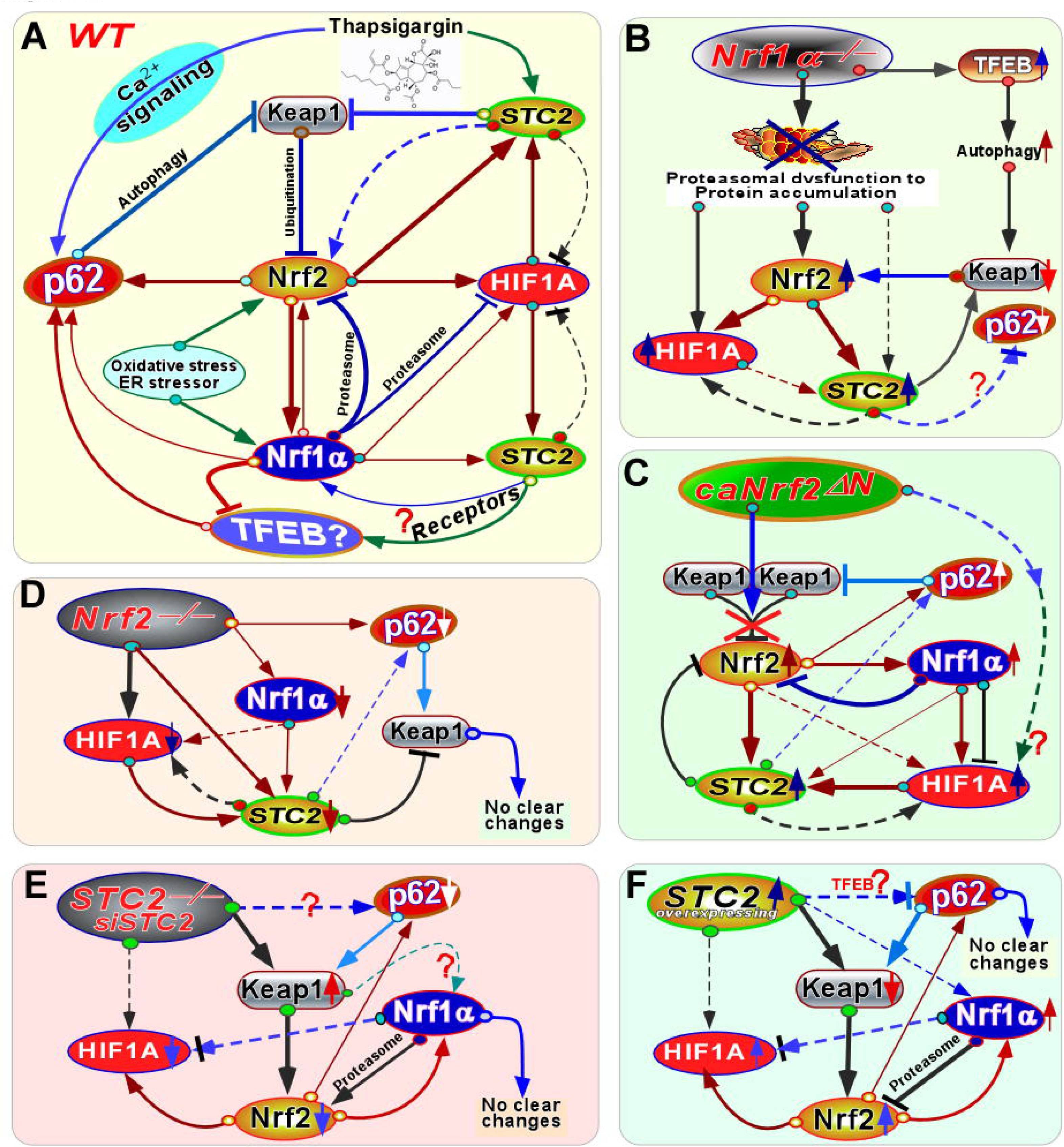
Distinct intrinsic status of key molecular-cellular regulatory models to provide a better understanding of distinction pathophysiological phenotypes. (A) A wild-type cellular-molecular inter-regulatory network is proposed for a better explanation of those key gene transcription (indicated by dark red lines) to core protein functions (illustrated by all other ways) in maintaining normal cellular homeostasis and even organ integrity. (B) The putative *Nrf1αNrf2α^-/-^*-specific cellular-molecular inter-regulatory network to give a better understanding of the mechanism dictating its unique pathological phenotype. (C) The caNrf2*^Δ^*^N^-led key molecular inter-regulatory network to explain its specific phenotype. (D) The Nrf2*^-/-^*-specific cellular-molecular inter-regulatory network to determine its phenotype. (E) A model proposed to explain the inter-regulatory network of between those core genes in *STC2^-/-^* cells. (F) The lentiv-STC2-leading molecular inter-regulatory network amongst those indicated key genes.

Conversely, STC2 had been also shown to interact with Nrf2 in mesenchymal stem cells [72]. In this study, our evidence has been presented revealing that the protein expression levels of Nrf2 was reduced by silencing of STC2 to antagonize Keap1 in *WT* and *Nrf1αNrf2α^-/-^* cell lines. This is also supported by further evidence that up-regulation of STC2 by TG was accompanied by significant down-regulation of Keap1 in WT, *Nrf1αNrf2α^-/-^*, and Nrf2 cell lines. Based on the fact that TG can also inhibit the transport of free Ca into the endoplasmic reticulum so as to increase the intracellular Ca^2+^ level and thus activate and/or prolong the Ca^2+^-mediated signaling pathways [54], it is inferable that STC2-triggered Ca^2+^ signaling may also play a role in the cytoprotective response to crosstalk with the Keap1-Nrf2 antioxidant pathway against various cellular stress (Figures 7C & 8A).

As such, the protein abundance of Nrf2 in caNrf2*^Δ^*^N^ cells (albeit its Keap1-binding domain lacks) remained to be down-regulation by silencing of STC2, as also accompanied by almost no changes in Keap1, implying an involvement of other targets except Keap1, such as Hrd1 or β-TrCP [73-75]. In addition, it should also be noted that the mRNA expression levels of Keap1 were largely unaltered in all examined cell lines (Figures S10F and S12), while its interactor p62 (called SQSTM1) at its mRNA levels was obviously down-regulated in *STC2^-/-^* and Nrf2*^-/-^* cell lines, albeit the upstream regulator TFEB is modestly up-regulated in both *Nrf1αNrf2α^-/-^* and lentiv-STC2 cell lines (Table S8). These suggest that STC2 is also likely involved in p62-mediated autophagy signaling (bi-directionally modulated by TFEB, Nrf1α and Nrf2) to monitor the abundance of Keap1. Moreover, modest up-regulation of Nrf1 by STC2 also occurred concomitantly with an exception of Keap1 and HIF1A that were marginally increased (Figure S10, F &G), but the detailed mechanism requires to be explored.

In further investigation of the biological role of STC2 in cell growth and proliferation, we have obtained a series of experimental evidence revealing that the clonogenicity of hepatoma and its cell-cycle turnover were evidently promoted by lentiv-STC2 overexpression, as well as the malignant growth of its xenograft tumors, but all opposite effects were manifested by knockout of *STC2^-/-^*. These demonstrate that STC2 is a potent tumor-promotor that plays a critical role in the progression of liver cancer. This is also supported by a previous study showing that ectopic STC2 expression markedly promoted hepatoma cell proliferation [29]. Altogether, these provide a clear explanation for discrepant phenotypes between *Nrf1αNrf2α^-/-^* and *^-/-^*-derived xenograft tumors. In *Nrf1αNrf2α^-/-^* cells, the increased expression of STC2 significantly promotes cell proliferation and in vivo malgrowth of its xenograft tumor. Conversely, such tumor malgrowth is almost completely abolished by loss of STC2, in line with the observation of nude mice inoculated Nrf2 cells owing to loss of its tumor-promoting function [76, 77]. In addition, significantly increased expression of STC2 can promote the pathogenic progression from steatosis to nonalcoholic steatohepatitis (NASH) [78], coincidently similar to the pathological phenotype of spontaneous NASH in liver-specific Nrf1*^-/-^* mice, along with its subsequent malignant transformation into hepatoma [7, 79].

In summary, this study provides a holistic perspective of the realistic scenario analysis integrated different sets of big data-mining with routine reductionist approaches, aiming to give a better understanding of the mechanisms underlying distinction pathophysiological phenotypes among all the examined *Nrf1αNrf2α^-/-^*, caNrf2, Nrf2, STC2, lentiv-STC2 cell lines, as compared with *WT* cells. Such distinct phenotypes should also be determined by different intrinsic status of a robust self-organized endogenous molecular-cellular network, with distinct feedback regulatory mechanisms (Figures 7C and 8). These selected stable states are predominantly dictated by altered programming from key gene transcription to mRNA translation into proteins at those core modular (and signaling) nodes of this network, along with their distinct topomorphisms shaped by a variety of post-transcriptional and post-translational modifications so to exert their different or even opposing functions in diverse subcellular topospatiotemporal self-organization systems. The overall homeodynamic states of such self-organizing systems are determined principally by their robustness and plasticity, i.e., two naturally-selecting but apparently-conflicting properties of the biological systems [80, 81], and can also be characterized by their nonlinear stochastic mathematic models [55-58]. Thereby, the robust homeostasis is successfully maintained by those evolutionally-conserved modular molecules and their interactive signaling pathways with distinct feedback regulatory mechanisms. The plasticity of the homeodynamic states are manifested primarily by a vast variety of adaptive responses to diverse cell stresses in different changing environments. From it, it is inferable that the existence of several nonlinear stochastic molecular events occurring at distinct strata (e.g., from gene transcription to mRNA translation into proteins) is uncovered by some seemingly-paradoxical data obtained from several databases and also in this study, to be presented in a holographic functional landscape as done as possible in realty. Such apparently-conflicting stochastic events are also likely triggered by potential double-edge effects of key modular molecules and their bi-directional feedback regulatory mechanisms. Altogether, all genetic and non-genetic drivers could be integrated as a selection force in Darwinanan dynamics to enable for a stochastic speciation of *Nrf1αNrf2α^-/-^*-deficient cells during carcinogenesis and ensuing cancer progression. Herein, our evidence demonstrates that significant upregulation of STC2 by hyper-expressed Nrf2, rather than its downstream HIF1A, in *Nrf1αNrf2α^-/-^* cells, as well in HCC tissues, leads to promotion of hepatoma cell proliferation and malgrowth of its xenograft tumor in nude mice. By contrast, upregulation of STC2 by HIF1A is also determined in caNrf2*^Δ^*^N^ cells. In turn, STC2 can also regulate Nrf2 by antagonizing its negative regulator Keap1, but conversely the latter Keap1 is also negatively regulated by Nrf2-target p62 so to form a dual feedback regulatory circuit.

However, loss of *STC2^-/-^* results in almost complete abolishment of both its deficient cell clonogenicity and xenograft tumor malgrowth, resembling the pathological phenotype of Nrf2*^-/-^*. Overall, this study highlights that like Nrf2, STC2 can serve as a potent tumor promotor, particularly in *Nrf1αNrf2α^-/-^*-deficient tumors, and may also be paved as a potential therapeutic target for relevant liver cancer.

## Supplemental Materials

The supporting information includes 15 supplemental figures and also eight supplemental tables.

## Acknowledgements

We are greatly thankful to all other members of Prof. Zhang’s laboratory (Chongqing University, China) for giving their invaluable help in this study. This work was funded by the National Natural Science Foundation of China (NSFC, 82073079, 81872336 and 91429305) awarded to Prof. Yiguo Zhang (at Chongqing University), and also was in part supported by the Initiative Foundation of Jiangjin Hospital affiliated to Chongqing University (2022qdjfxm001).

## Author Contributions

Both Q.B. and Y.D designed and performed most of the experiments, and wrote the manuscript draft. Q.W. and R.D. participated in bioinformatic analysis. S.H. provided critical suggestions for this work. Z.P. did pathohistological analysis. Lastly, Y.Z. designed and supervised this study, parsed all the data, helped to prepare all the figures, wrote and revised this manuscript. All authors have read and approved this version of the manuscript for publication.

## Data availability statement

The datasets analyzed for this study can be found in online repositories. The names of the repository/repositories and accession number(s) can be found in the article, along within Supplementary Material.

## Ethics statement

The animal study was reviewed and approved by the University Laboratory Animal Welfare and Ethics Committee (with two institutional licenses SCXK-PLA-20120011 and SYXK-PLA-20120031).

---

**Fig. S1. Identification of *Nrf1αNrf2α^-/-^*, Nrf2*^-/-^*, *Δ*N caNrf2 cell lines and STC2-specific antibody.**

(A) The protein expression abundances of Nrf1α and Nrf2 in *WT* HepG2 cells and its derivative*-/--/-*and caNrf2 cell lines were determined by Western blotting. Nrf1α, Nrf2

B) *WT* cells were transfected with a STC2 expression construct or empty vector and then subjected to Western blotting to verify the accuracy of STC2-specific antibody and its V5-tagged proteins.

(C) *WT* cells were treated or not treated with (1μM dose) TG to induce the STC2 expression and subjected to Western blotting to verify the accuracy of STC2-specific antibody.

**Fig. S2. Establishment of STC2 and *^-/-^* STC2 cell lines.**

(A) The genomic DNA sequencing to identify two mutants of STC2 in selected cell lines, which is thus designated as STC2^insC^ and *STC2^-/-^* cell lines, respectively.

(B) The mutagenesis mapping of ATGs (as putative translation start codons at #1, #2 and #3 positions mutated into CTGs) within the open reading frame of STC2.

(C) HepG2 cells were transfected with the above-described STC2 mutants #1, #2 and #3, and then subjected to Western blotting analysis of the STC2 protein expression levels.

**Fig. S3. The enrichment analysis of DEGs in liver cancer tissues compared to normal liver tissues.**

(A, B) Two VENN maps of DEGs significantly up-regulated or down-regulated in liver cancer tissues as compared to the normal liver tissues, which were obtained by distinct analysis packages DESeq2, LIMMA and edgeR.

(C to E) Those up-regulated DEGs enriched respectively in the GO cell components, biological processes and the KEGG pathways in liver cancer tissues when compared with the normal tissues.

(F to H) Those down-regulated DEGs enriched respectively in the GO cell components, biological processes and the KEGG pathways in liver cancer tissues when compared with the normal tissues.

**Fig. S4. Analysis of liver cancer data obtained from TCGA.**

(A) Principal component analysis of HCC samples obtained from the TCGA database. (B) A volcano map of DEGs in HCC analyzed by the DESeq2 package.

(C) A heat-map of the expression values of 30 top DEGs significantly in HCC.

D) The impact of STC2, CBX2, ADAM1 or AKR1D1 on the overall survival rate of HCC patients was evaluated by the Kaplan-Meier’s method.

**Fig. S5. The DEGs in TCGA-LIHC tissues and *Nrf1αNrf2α^-/-^* or *^-/-^* cell lines as compared with their controls.**

(A to C) Three VENN maps of DEGs in the TCGA-LIHC tissues intersected with other DEGs in *Nrf1αNrf2α^-/-^*, *^-/-^* or caNrf2 cell lines selected by comparison with their *WT* counterparts.

(D) The FPKM values of DEGs in *WT* and *^-/-^* cell lines, all of which are up-regulated in LIHC and *^-/-^* cells.

(E) The FPKM values of DEGs in *WT* and *^-/-^* cell lines, all that are down-regulated in LIHC and *^-/-^* cells.

(F) The FPKM values of DEGs in *WT* and *-/-*ll lines, all of which are up-regulated in LIHC but down-regulated in *Nrf1αNrf2α^-/-^* cells. Nrf1αce

G) The FPKM values of DEGs in *WT* and *Nrf1αNrf2α^-/-^* cell lines, that are all down-regulated in LIHC but also up-regulated in *Nrf1αNrf2α^-/-^* cells.

(H) The FPKM values of DEGs in *WT* and Nrf2*^-/-^* cell lines, that are all up-regulated in LIHC and Nrf2*^-/-^* cells.

(I) The FPKM values of DEGs in *WT* and Nrf2*^-/-^* cell lines, that are all up-regulated in LIHC but also down-regulated Nrf2 cells.

(J) The FPKM values of DEGs in *WT* and *^-/-^* cell lines, that are all down-regulated in LIHC, but also down- or up-regulated in *^-/-^* cells respectively.

**Fig. S6. Comparative analysis of LIHC data in TCGA and transcriptome data of *WT* and*^Δ^*^N^ cell lines. caNrf2**

(A) The FPKM values of DEGs selected in *WT* and caNrf2*^Δ^*^N^ cell lines, that are all up-regulated in LIHC but also down-regulated in caNrf2*^Δ^*^N^ cells.

(B) The FPKM values of DEGs in *WT* and *^Δ^*^N^ cell lines, that are all up-regulated in LIHC and *^Δ^*^N^ cells.

(C) The FPKM values of DEGs in *WT* and *^Δ^*^N^ cell lines, that are all down-regulated in LIHC and *^Δ^*^N^ cells.

(D) The FPKM values of DEGs in *WT* and *^Δ^*^N^ cell lines, that are all down-regulated in LIHC but up-regulated in caNrf2*^Δ^*^N^ cells.

(E) The expression levels of CBX2 in LIHC obtained from the Ualcan database and its effect on the survival of HCC patients evaluated by the Kaplan-Meier Plotter database.

(F) The expression levels of HOXD9 in LIHC obtained from the Ualcan database and its effect on the survival of HCC patients evaluated by the Kaplan-Meier Plotter database.

**Fig. S7. The correlation between those key gene expression levels in the liver cancer database.**

(A, B) The correlation between the expression levels of Nrf1 and GCLM, PSMB7 in the LIHC database.

(C, D) The correlation between the expression levels of Nrf2 and GCLM, Nrf1 in the LIHC database.

(E to H) The correlation between the expression levels of Nrf1, Nrf2, HIF1A, AHR and STC2 in the LIHC database.

**Fig. S8. The ChIP-Sequencing analysis of Nrf1 and Nrf2 on the Encode database.**

(A) Nrf1 binds to the promoter regions of *GCLM* or STC2 in HepG2 cells (data obtained from the Encode database).

(B) Nrf2 binds to the promoter regions of *GCLM* or STC2 in HepG2 cells (data obtained from the Encode database).

**Fig. S9. Analysis of transcription factors binding to the promoter region of genes.**

(A) Nrf1 and Nrf2 bind to the promoter region of *HIFA* in HepG2 cells (data obtained from the Encode database).

(B) Distinct effects of Nrf1, Nrf2 and HIF1A on distinct lengths of the STC2 promoter were detected by their relevant luciferase reporter genes that are co-transfected into HepG2 cells. The resulting data are shown graphically (n = 3 x 3, $ p < 0.05, $$ p < 0.01, NS = no statistical difference).

**Fig. S10. Analysis of gene expression changes in HepG2 cells after knockout or overexpression STC2.**

(A) The correlative heat-map of samples employed for transcriptome sequencing.

(B) A boxplot of their expression quantification distribution in the examined samples.

(C to E) Three volcano maps of DEGs in *STC2^-/-^* vs WT, Lentiv-STC2 vs *WT* or respectively. STC2 cell lines are illustrated,

(F) The TPM values of those indicated genes, including *STC2, Nrf1, Nrf2, HO-1, HO-2, NQO1, GCLC, GCLM, SQSTM1, HIF1A, HIF1AN, HILPAD, SLC2A1, VEGFA and Keap1* in *WT*, *STC2^-/-^* and Lentiv-STC2 cells.

(G) The *WT* HepG2 cells were transfected with a STC2-expressing plasmid and then subjected to Western blotting analysis of the protein expression changes of STC2, Nrf1, HIF1A and Keap1.

**Fig. S11. The enrichment analysis of significantly DEGs in*-/-* STC2 or Lentiv-STC2 cells by GO and KEGG methods.**

(A) The gene ontology (GO) enrichment analysis of DEGs for cell components, biological processes, and molecular functions in *STC2^-/-^* (left panel) or Lentiv-STC2 (right panel) vs *WT* cell lines. The top 20 highly representative GO terms are shown to be classified in DEGs.

(B) The KEGG pathway enrichment of DEGs between *STC2^-/-^* or Lentiv-STC2 vs *WT* cell lines. The graph shows the top 20 significantly enriched KEGG pathways.

**Fig. S12. The FPKM values of key genes in WT, *Nrf1αNrf2α^-/-^*, Nrf2*^-/-^* and caNrf2 cell lines, as they were deciphered*-/-*STC2 or Lentiv-STC2 cell lines when compared with *WT* controls (Figure 6D).**

**Fig. S13. The changes of those DEGs governing critical transcription factors regulated by Nrf1, Nrf2 and STC2 in Lentiv-STC2, *Nrf1αNrf2α^-/-^*, caNrf2*^Δ^*^N^ and*-/-*WT controls. Nrf2 cell lines when compared with**

**Fig. S14. The changes of those DEGs governing critical transcription factors regulated by Nrf1, Nrf2 and STC2 in *STC2^-/-^-/- Δ*N *-/-*, Nrf1α, caNrf2 and Nrf2 cell lines when compared with *WT* controls.**

**Fig. S15. The changes of those DEGs possibly involved in the Ca^2+^-relevant pathways in *Nrf1αNrf2α^-/-^*, Nrf2*^-/-^*, *Δ*N *-/-* caNrf2, Lentiv-STC2 and STC2 cell lines as compared with *WT* controls.**

