## Supplementary figures and images for "Stanniocalcin 2 (STC2) is a potent biomarker of hepatocellular carcinoma with its expression being augmented in Nrf1α-deficient cells, but diminished in Nrf2-deficient cells"

### FigureS1.tif

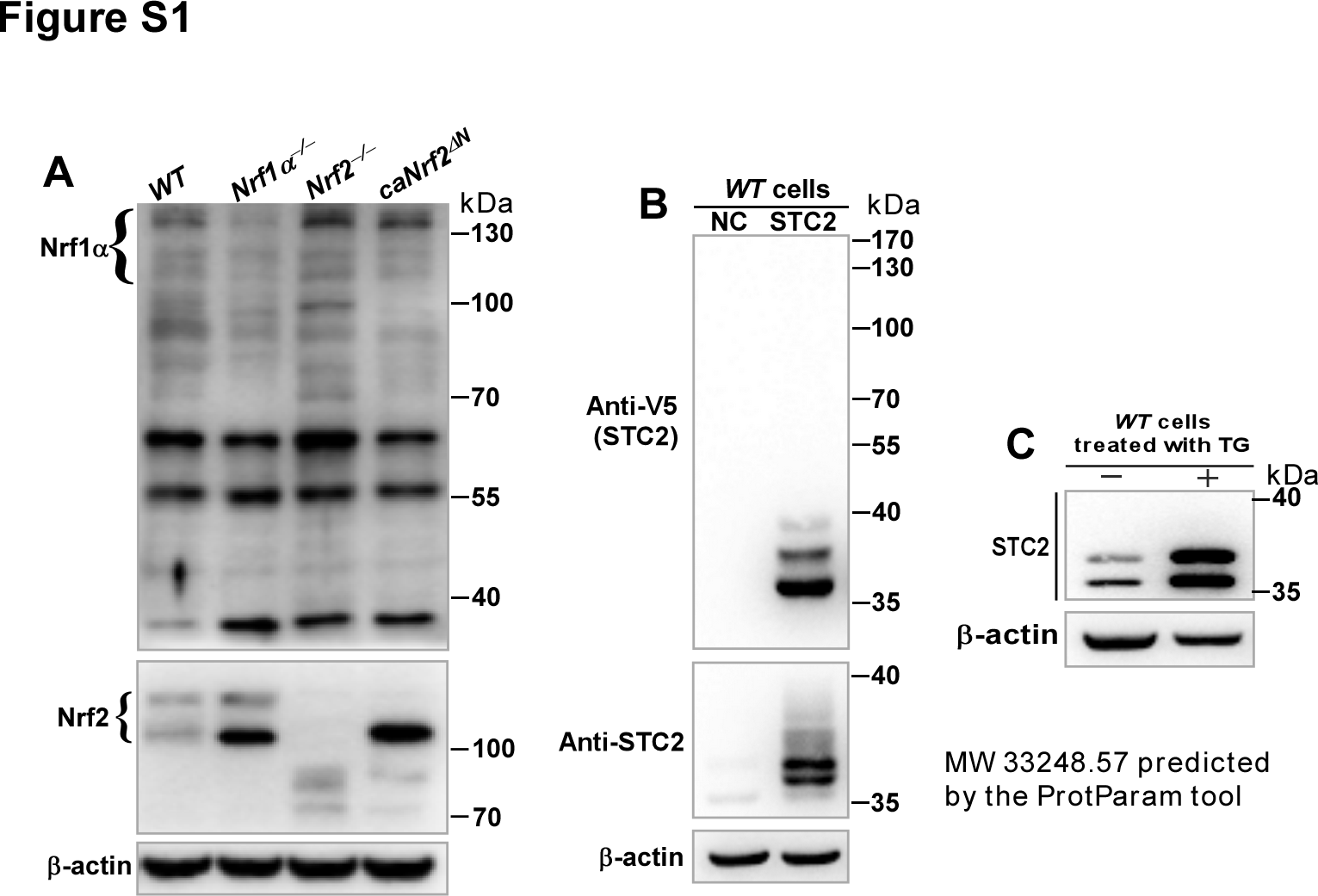

### FigureS2.tif

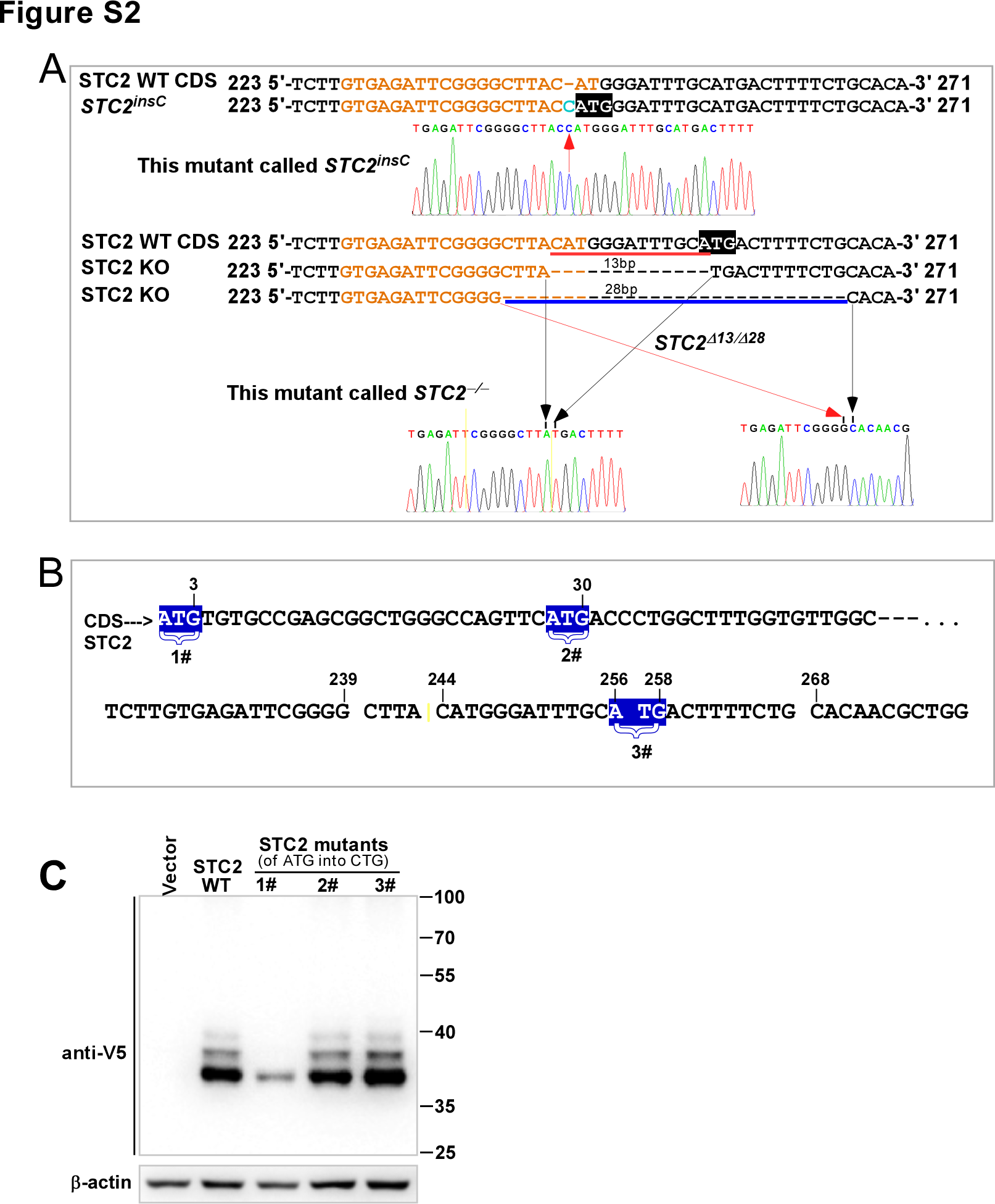

### FigureS3.tif

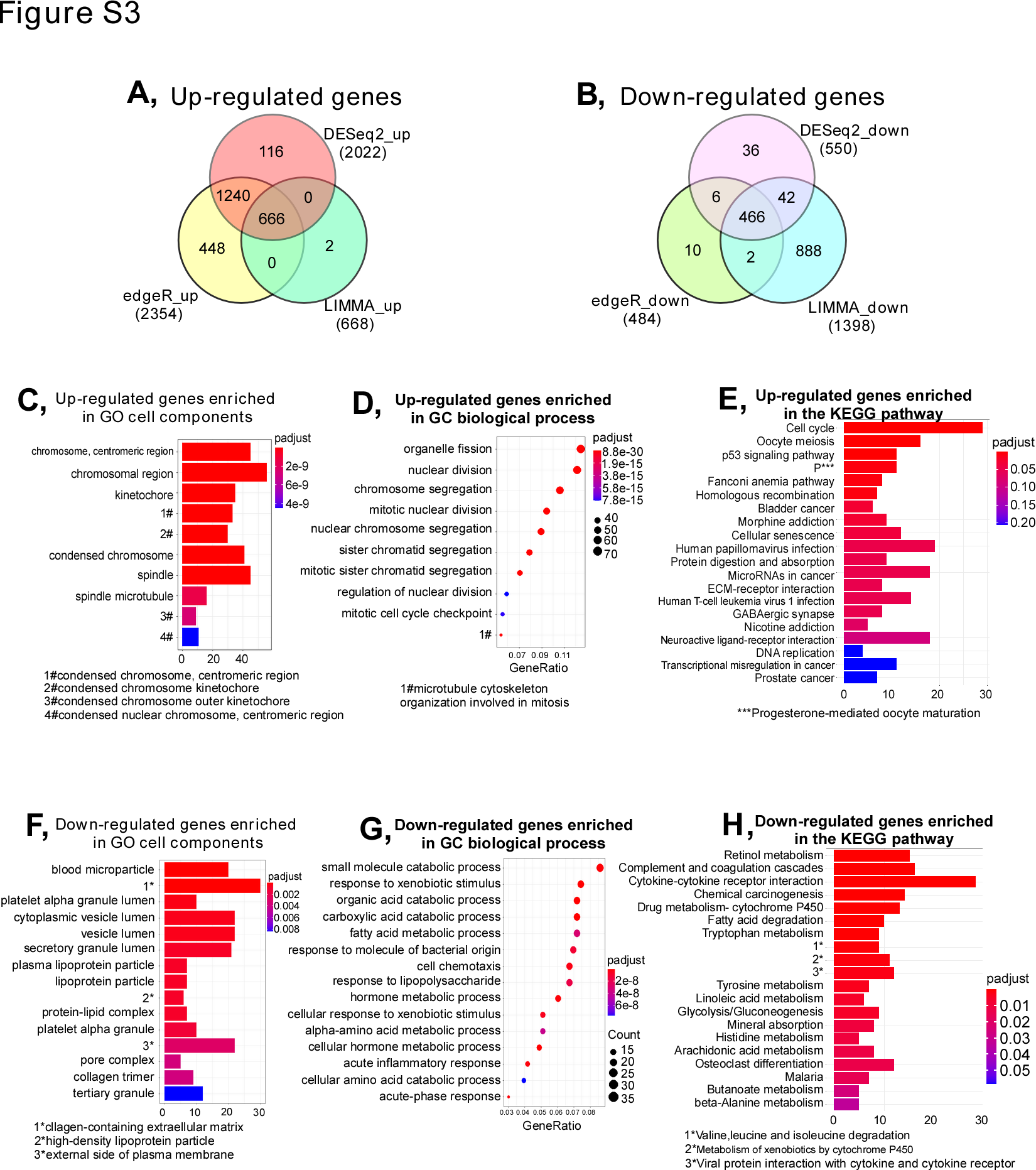

### FigureS4.tif

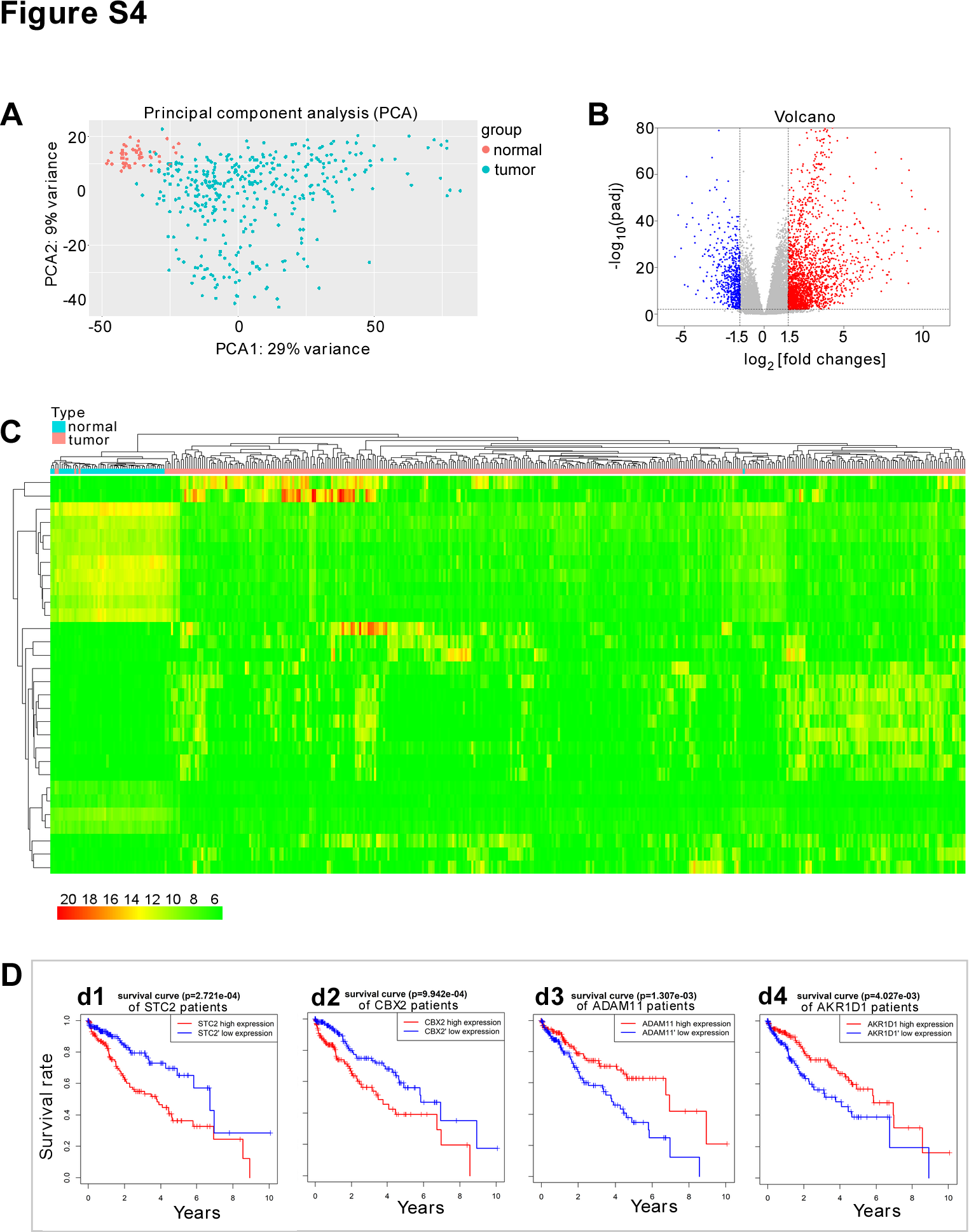

### FigureS5.tif

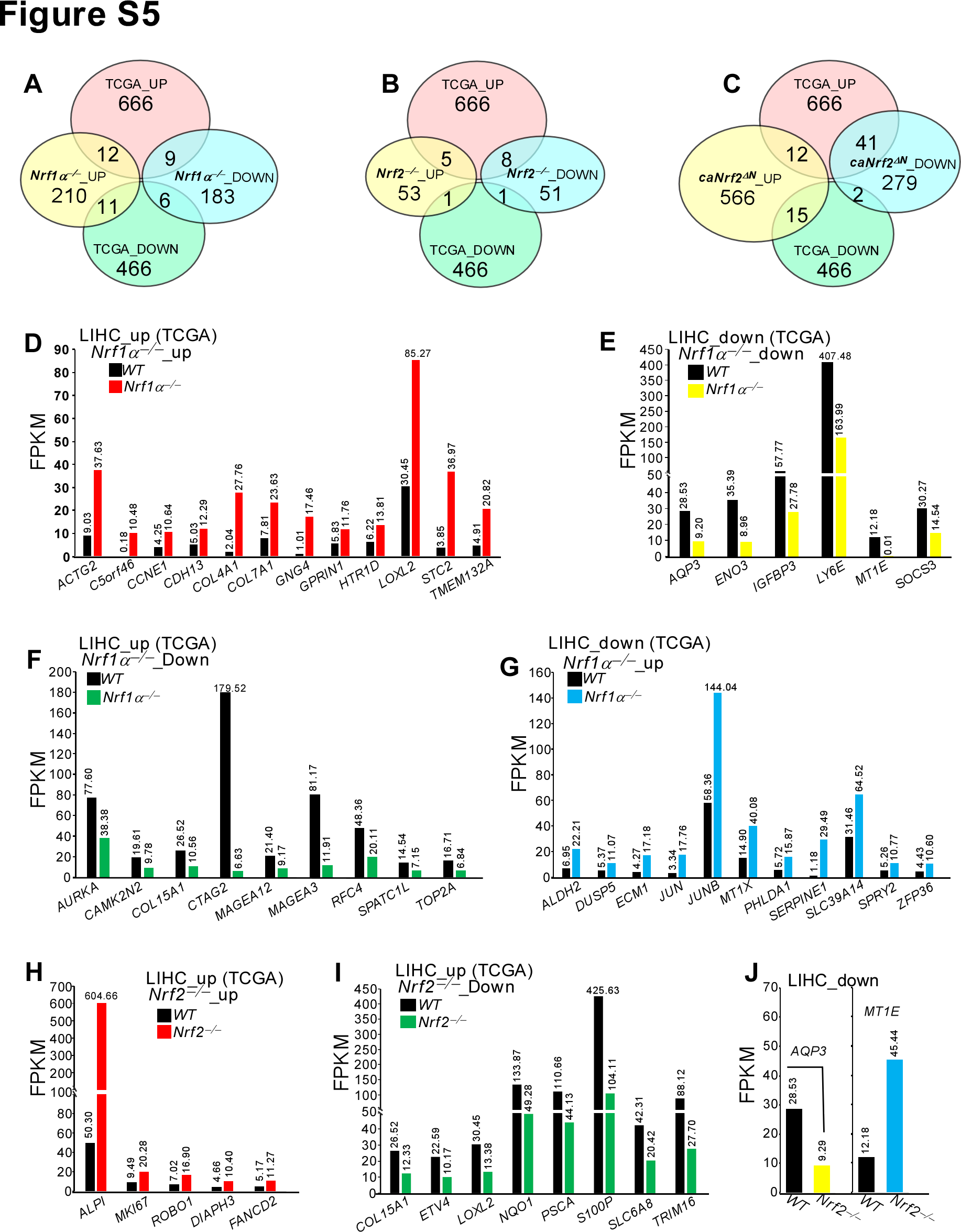

### FigureS6.tif

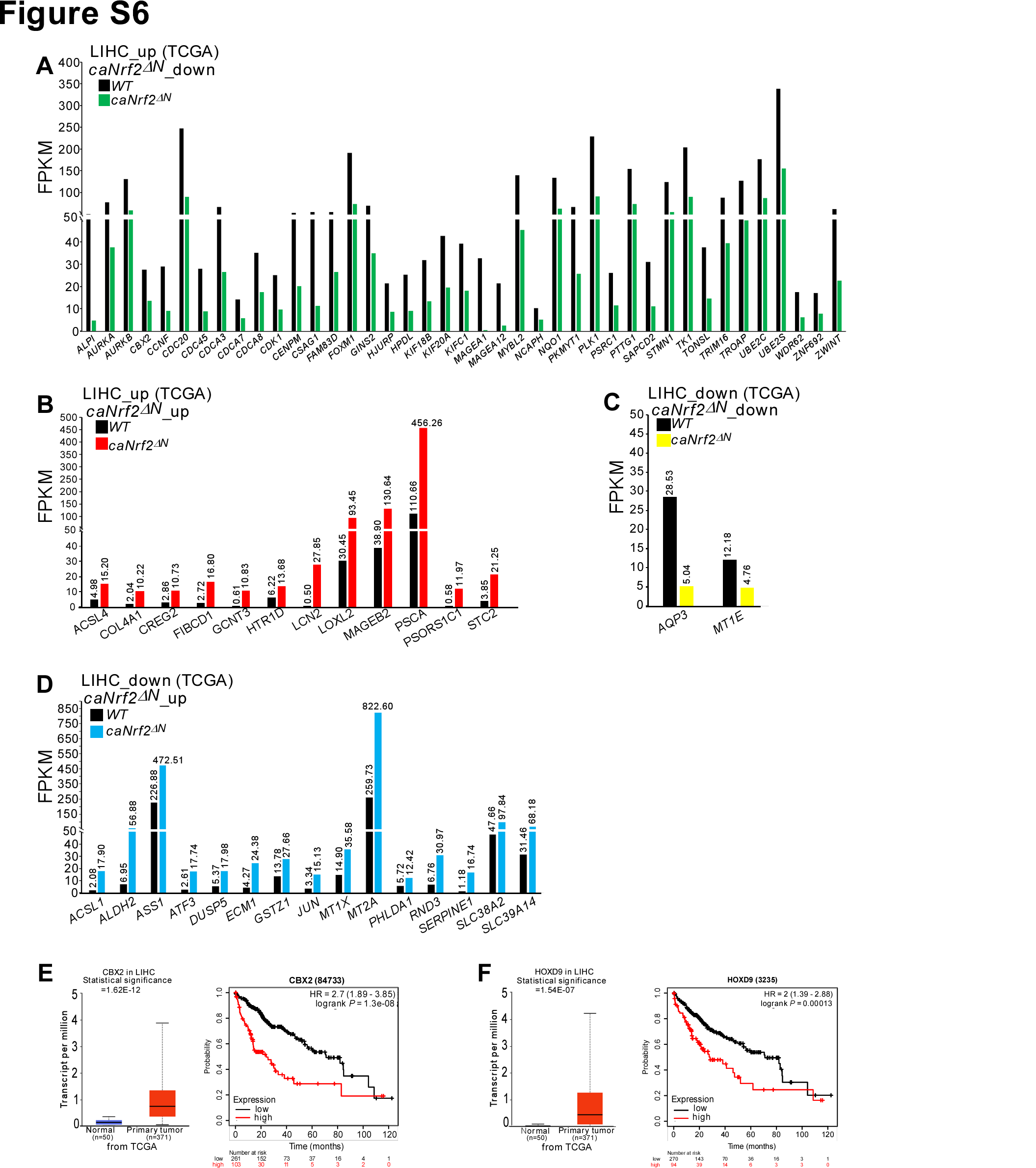

### FigureS7.tif

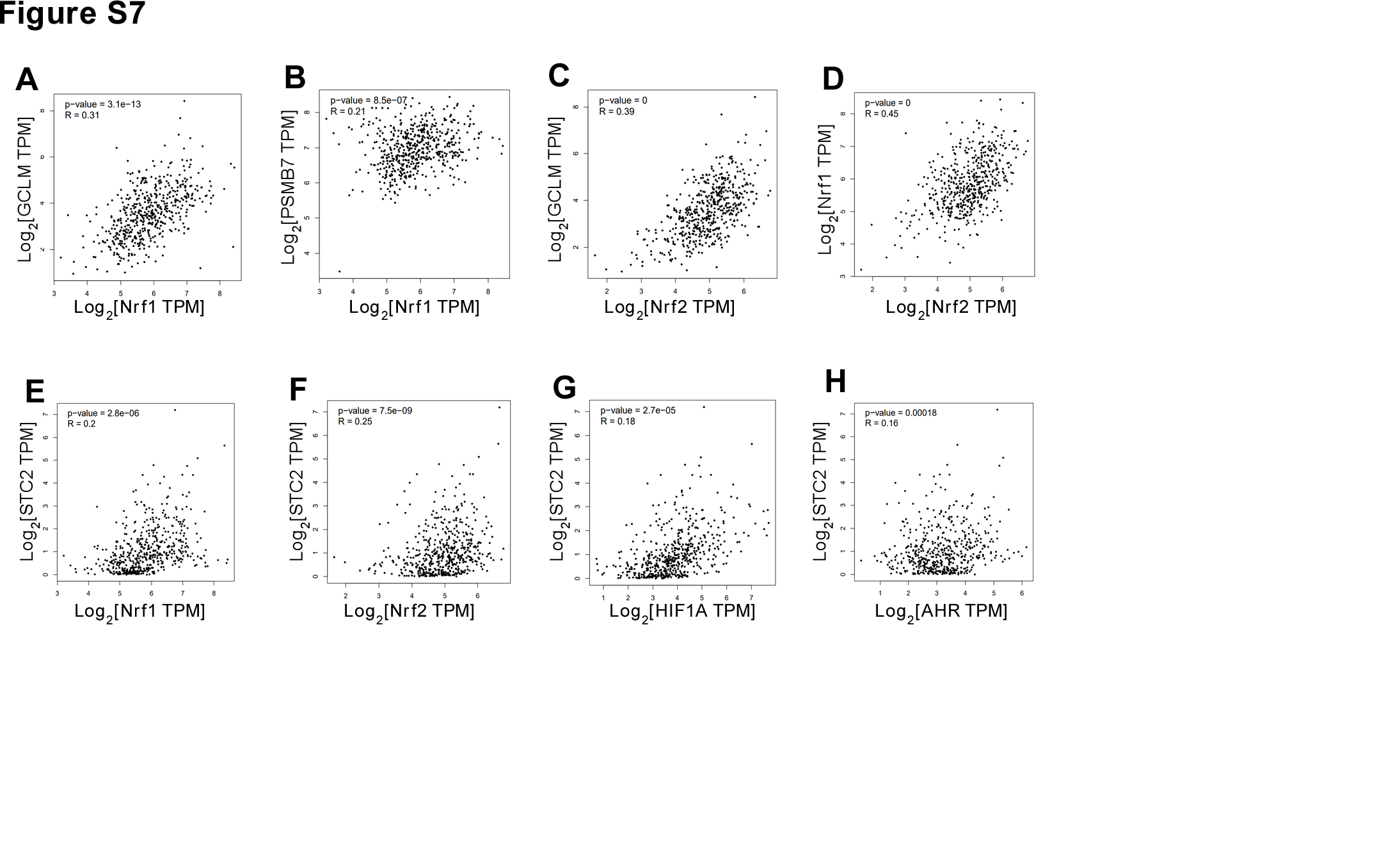

### FigureS8.tif

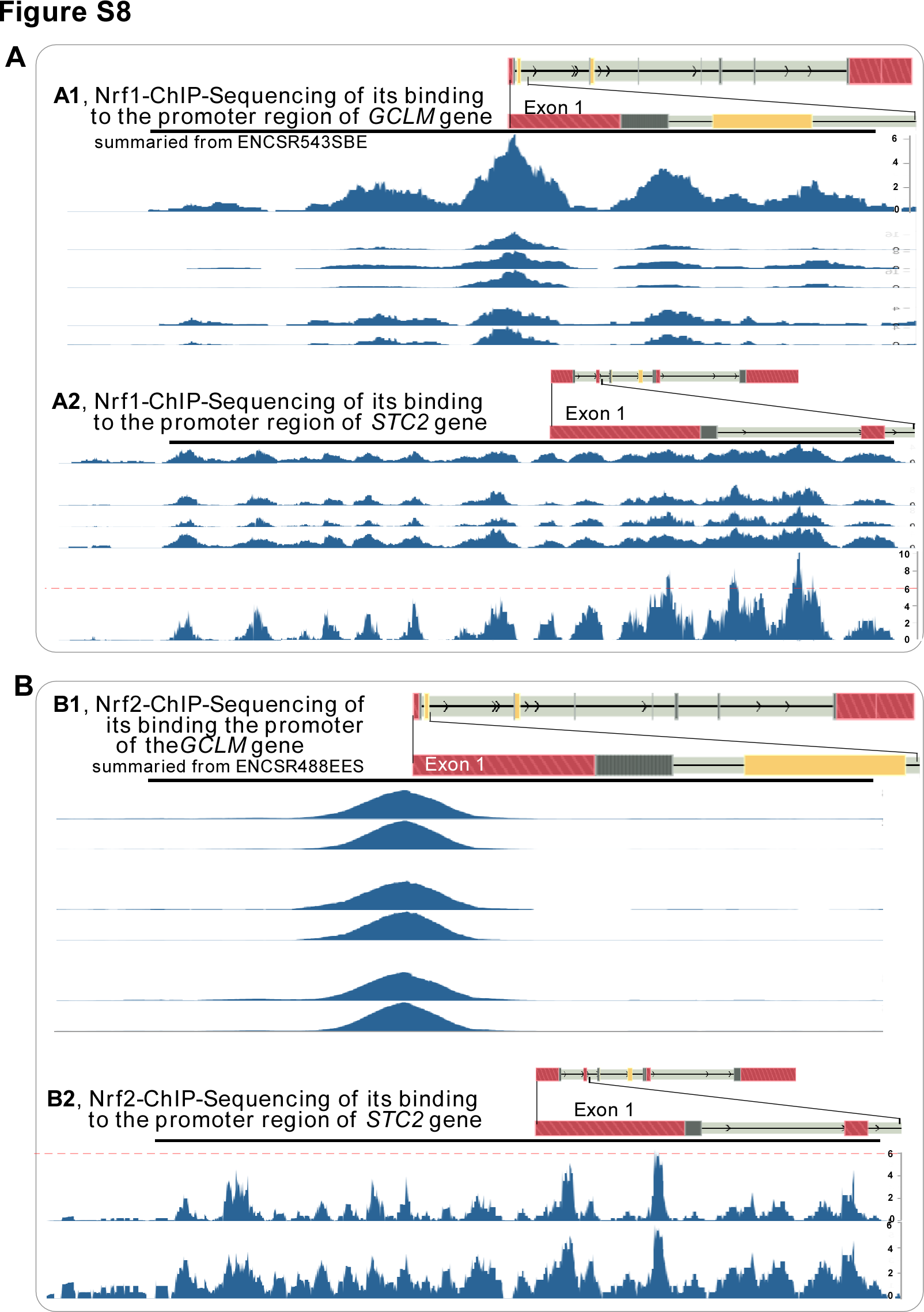

### FigureS9.tif

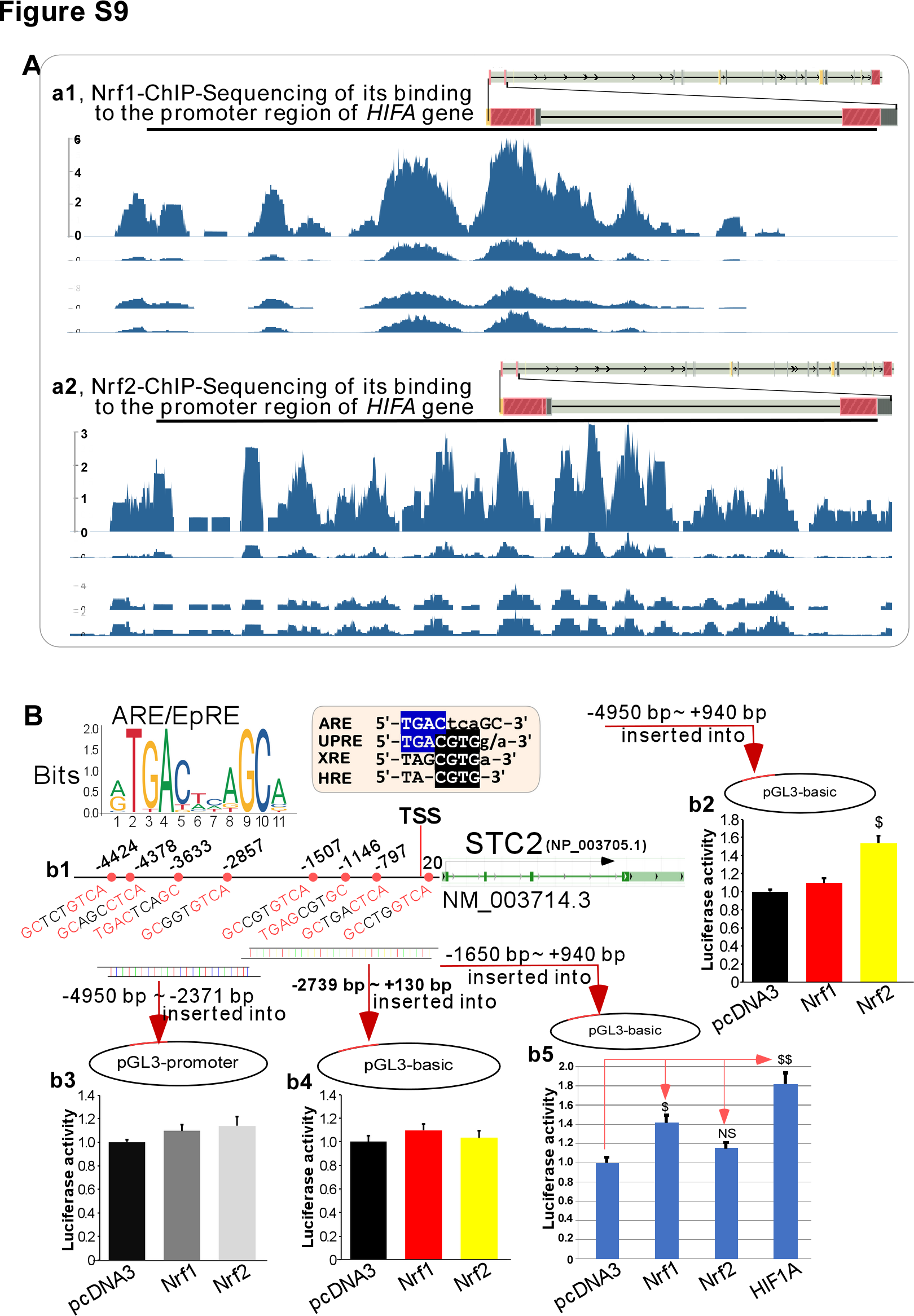

### FigureS10.tif

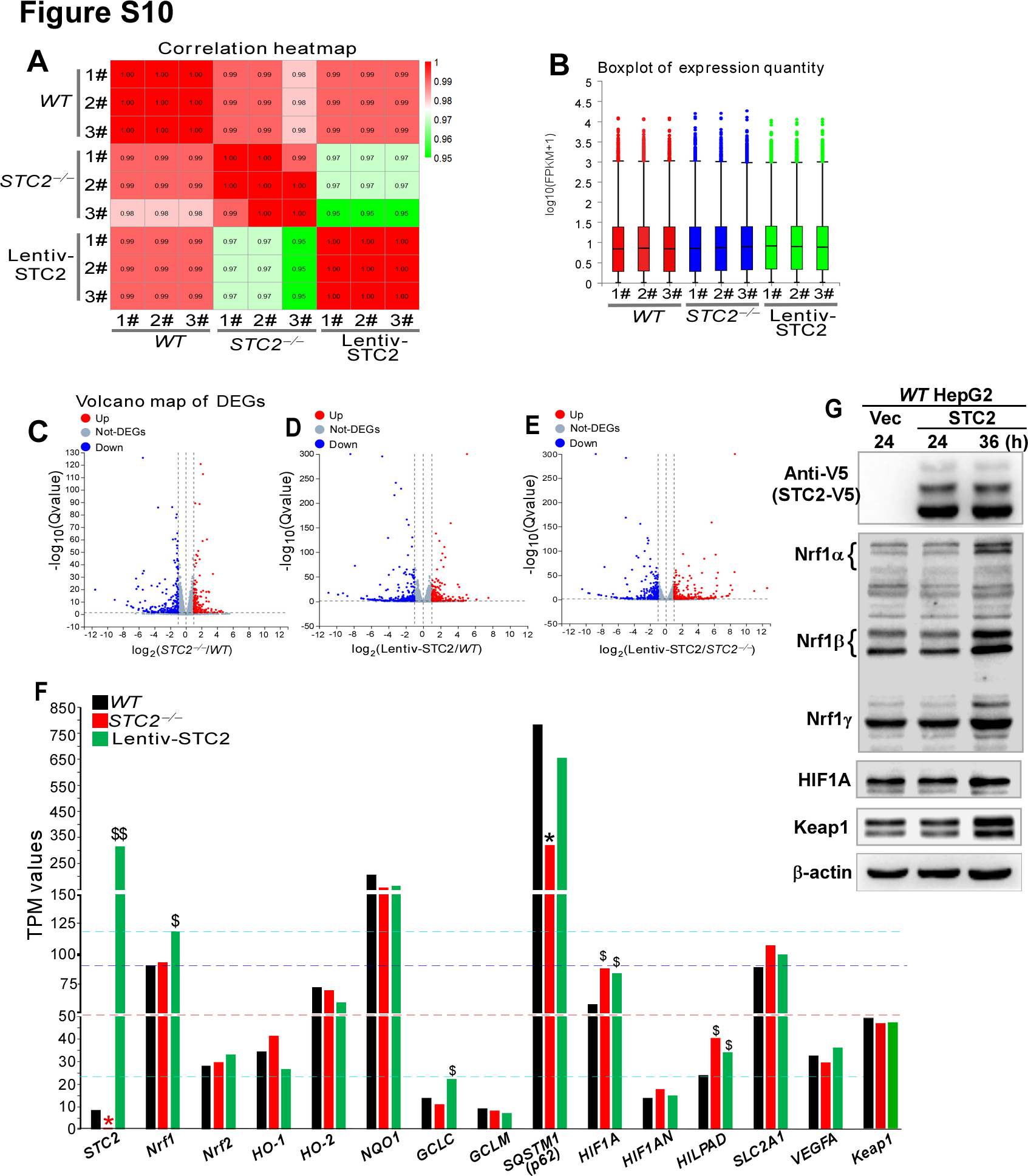

### FigureS11.tif

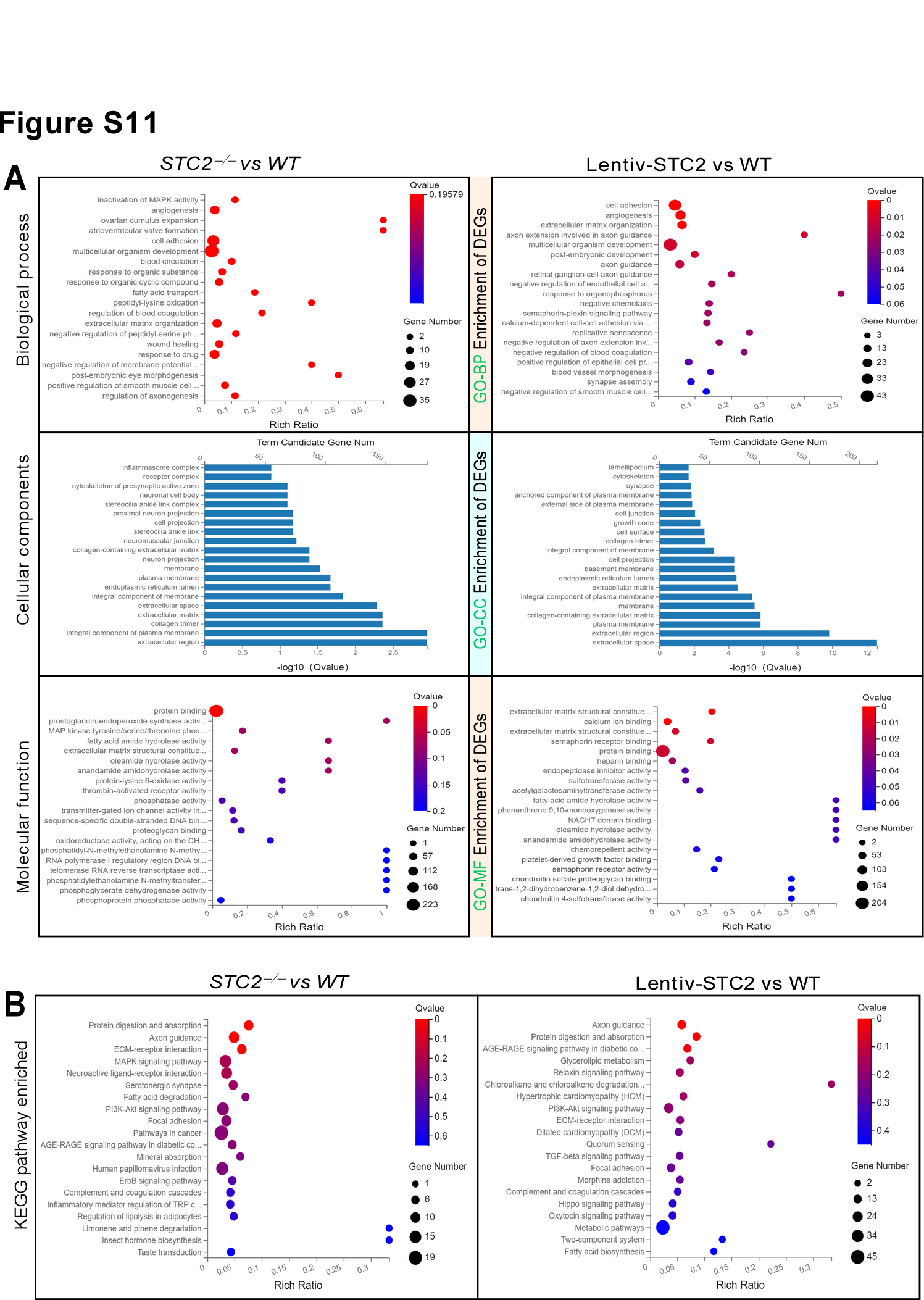

### FigureS12.tif

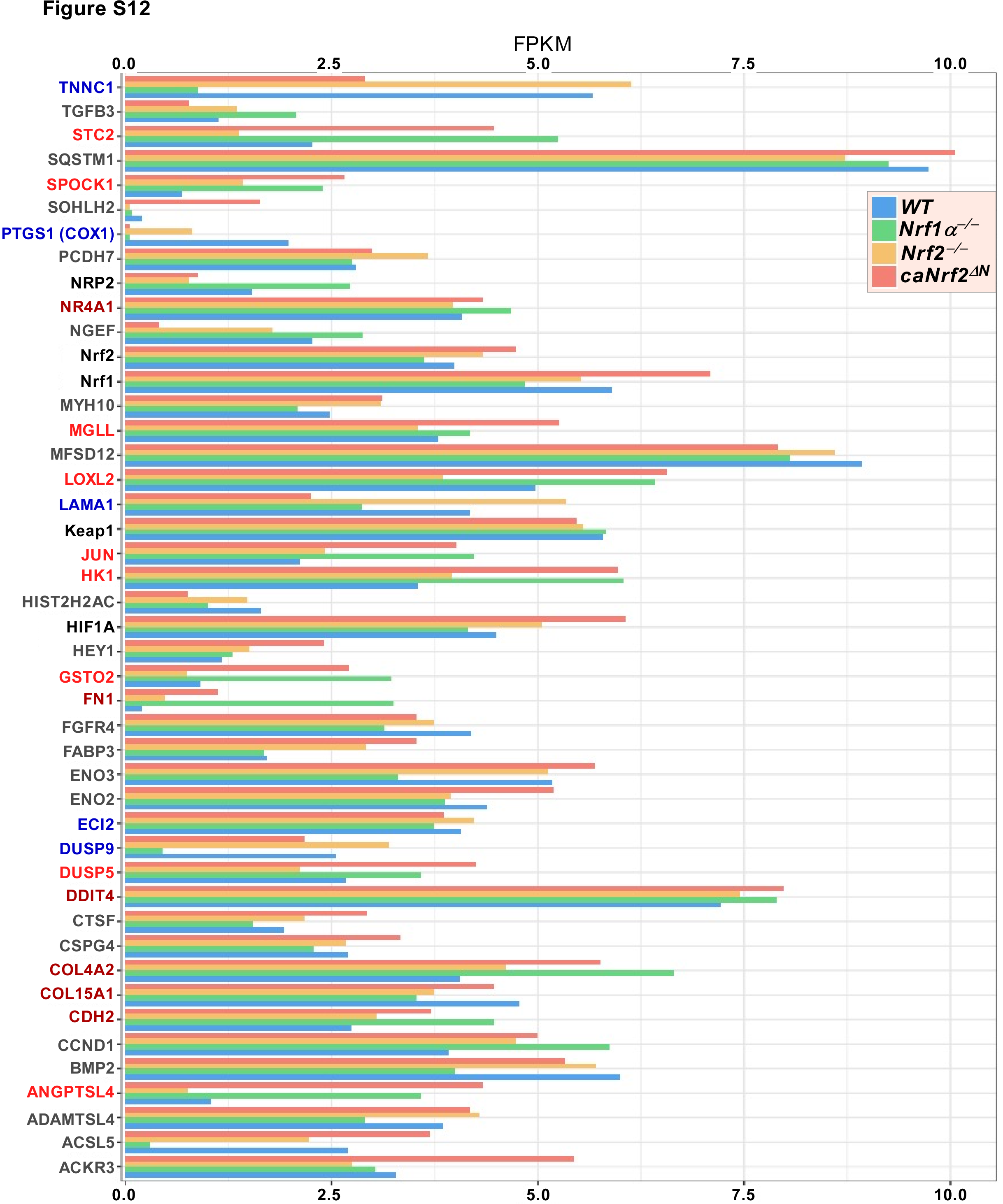

### FigureS13.tif

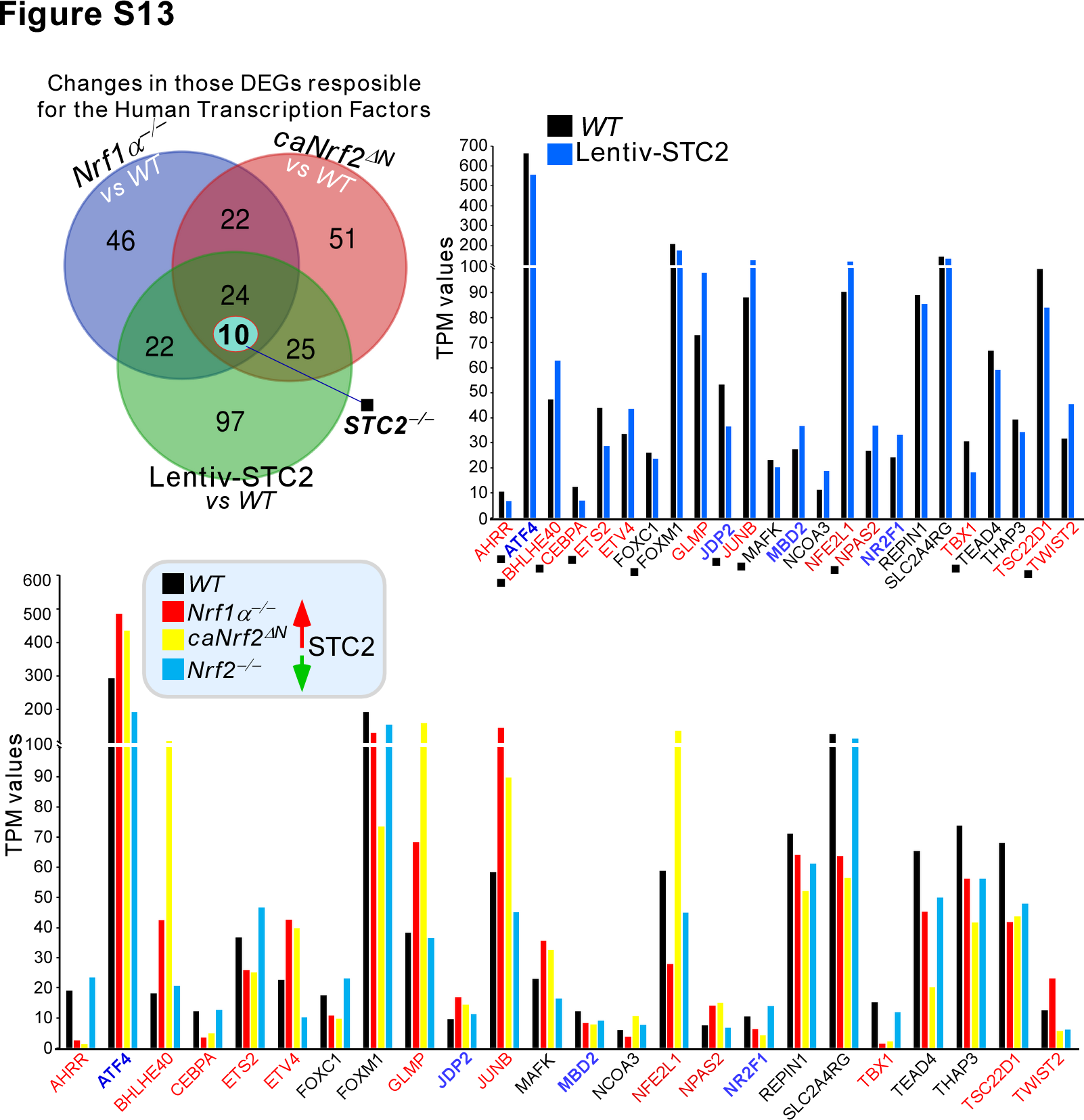

### FigureS14.tif

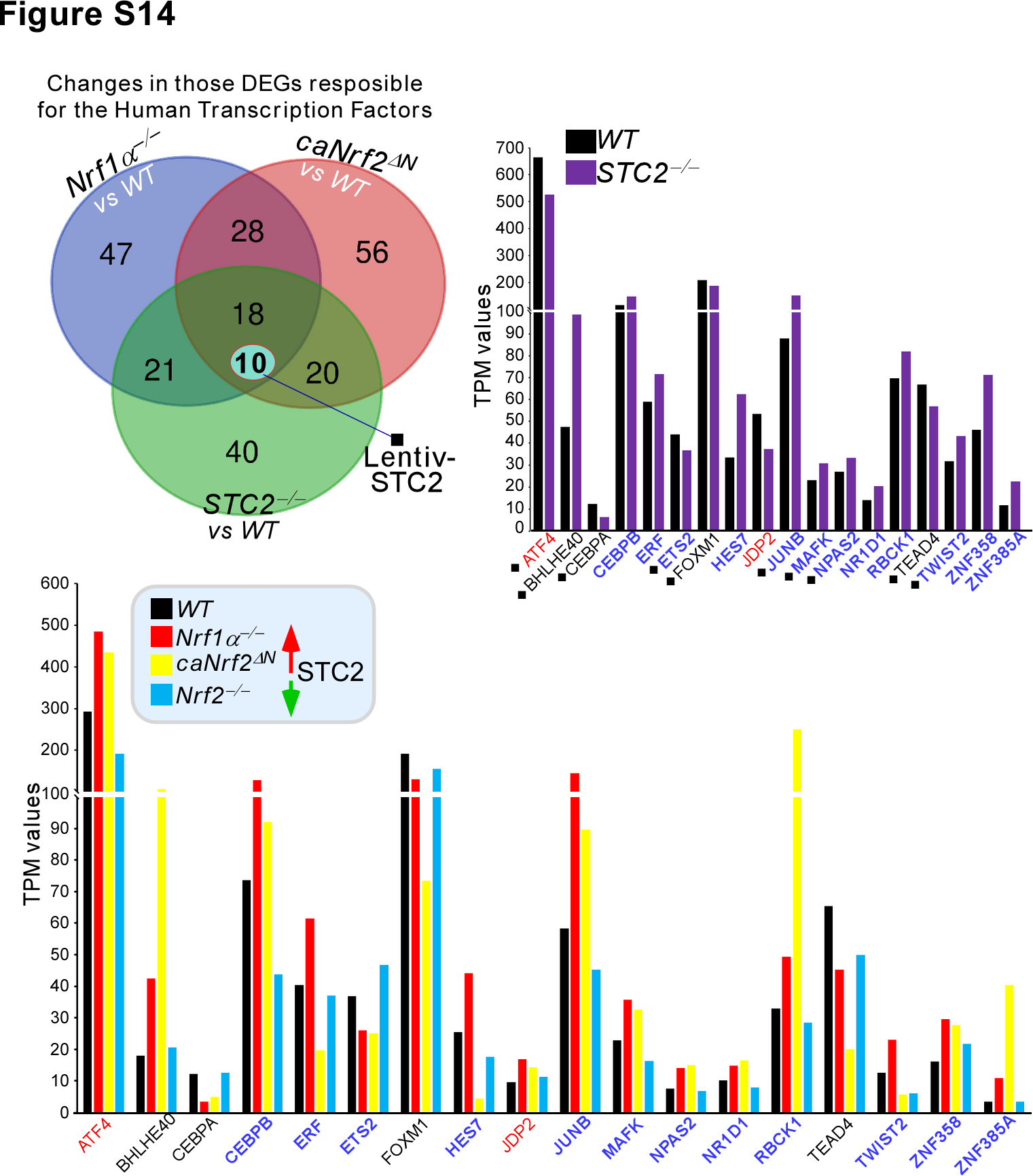

### FigureS15.tif

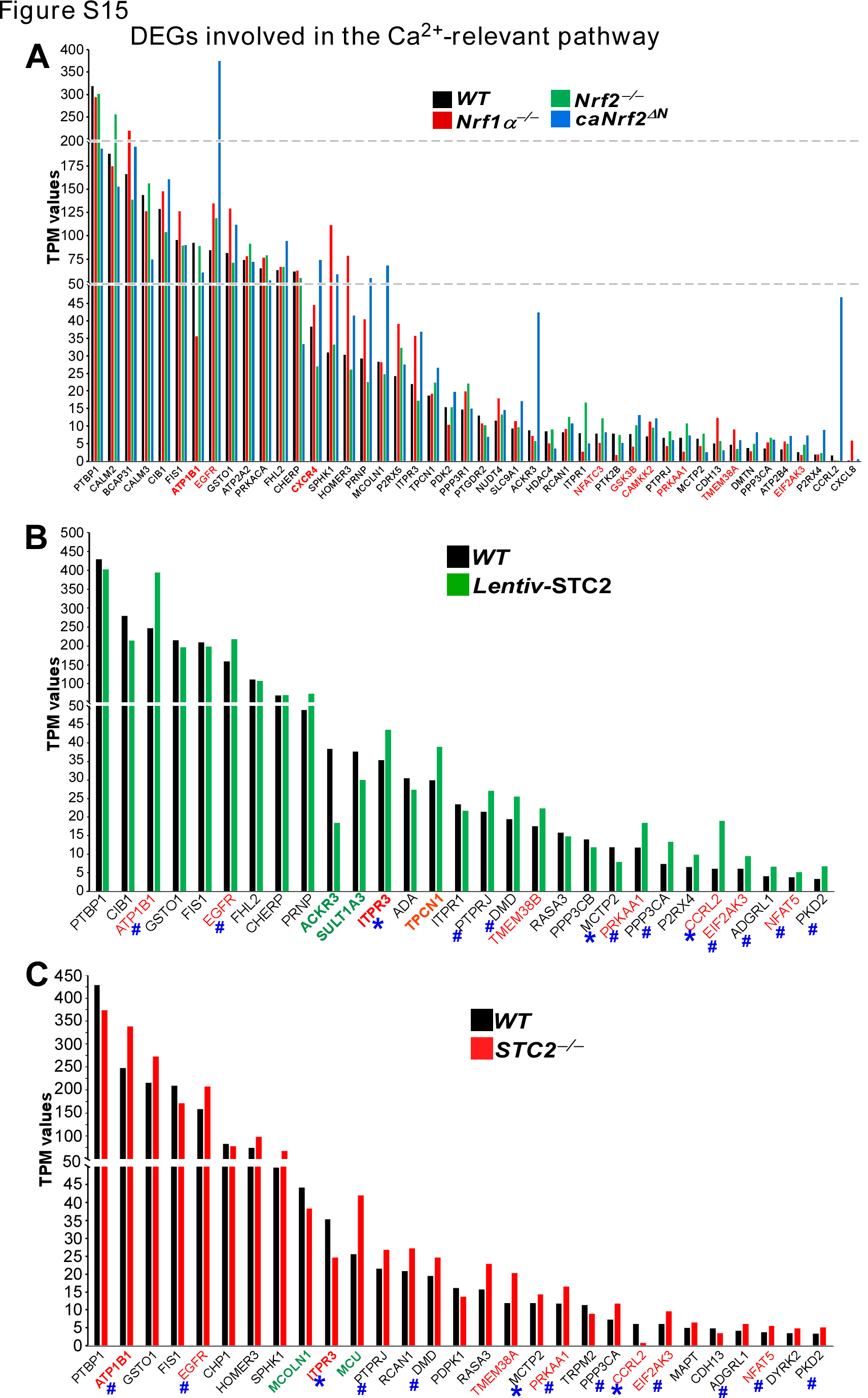
