## Supplementary material for "Stanniocalcin 2 (STC2) is a potent biomarker of hepatocellular carcinoma with its expression being augmented in Nrf1α-deficient cells, but diminished in Nrf2-deficient cells": Table S1-8

**Table S1**. Three pairs of siRNA sequences used in this study

| **siRNA** | **Nucleotide sequence (5′→ 3′)** |
| --- | --- |
| siNrf2 | Sense: GUUUGGGAGGAGCUAUUAUdTdT |
| Antisense: AUAAUAGCUCCUCCCAAACdTdT | |
| siHIF1A | Sense: CUGAUGACCAGCAACUUGAdTdT |
| Antisence: UCAAGUUGCUGGUCAUCAGdTdT | |
| siSTC2 | Sense: GUGGAGAUGAUCCAUUUCAdTdT |
| Antisense: UGAAAUGGAUCAUCUCCACdTdT | |

**Table S2**. The primer pairs used for quantitative RT-PCR

| **Genes** | **Nucleotide sequences (5′→ 3′)** |
| --- | --- |
| Actin | FP: GCGGCACCACCATGTACCCTG |
|  | RP: TCCACACGGAGTACTTGCGCTCA |
| Nrf2 | FP: TGTGGCATCACCAGAACACTC |
|  | RP: TCCAGGGGCACTATCTAGCTC |
| Nrf1 | FP: TGAAGCCCACCAAGACCGAAA |
| STC2 | RP: GCCTCTTCCTGTACACTGACC  FP: GCCTGTGCTCCATCTTG  RP: TTTGGGTGGCTCTTGC |
| HMOX1 | FP: CCAGGCAGAGAATGCTGAGTTC |
|  | RP: AAGACTGGGCTCTCCTTGTTGC |
| HIF1A | FP: TATGAGCCAGAAGAACTTTTAGGC |
|  | RP: CACCTCTTTTGGCAAGCATCCTG |
| GLUT1 | FP: CTGCAACGGCTTAGACTTCGAC |
|  | RP: TCTCTGGGTAACAGGGATCAAACA |
| VEGFA | FP: TTGCCTTGCTGCTCTACCTCCA |
|  | RP: GATGGCAGTAGCTGCGCTGATA |
| HILPDA | FP: AAGCATGTGTTGAACCTCTACC |
|  | RP: TGTGTTGGCTAGTTGGCTTCT |

**Table S3**. The gRNA sequences for STC2 knock out

| **Targets** | **gRNA sequence (5′→ 3′)** |
| --- | --- |
| STC2-1 | FP: AAACACCGCCTTTGACCCGGCGCGGGGG |
|  | RP: CTCTAAAACCCCCCGCGCCGGGTCAAAGG |
| STC2-2 | FP: AAACACCGGTGAGATTCGGGGCTTACAT |
|  | RP: CTCTAAAACATGTAAGCCCCGAATCTCAC |

**Table S4**. The eight-gene-based prognostic model for HCC established on the COX base

| **Gene** | **Coef** | **HR** | **Se-Coef** | **Z value** | ***p*_value** |
| --- | --- | --- | --- | --- | --- |
| GAGE2A | 0.360198373 | 1.433613776 | 0.160782263 | 2.240286747 | 0.025072313 |
| **STC2** | 0.188216476 | 1.207094794 | 0.098265155 | 1.915393873 | 0.055442292 |
| CBX2 | 0.259867475 | 1.296758222 | 0.116726179 | 2.226299847 | 0.025994104 |
| SPP1 | 0.084721269 | 1.08841365 | 0.037326635 | 2.26972697 | 0.023224154 |
| HOXD9 | 0.243178713 | 1.275296516 | 0.108604215 | 2.239127756 | 0.025147605 |
| PYDC1 | 0.556684801 | 1.744878282 | 0.208241245 | 2.673268692 | 0.007511606 |
| TKTL1 | 0.204251776 | 1.226606945 | 0.107439548 | 1.901085588 | 0.057290803 |
| ZDHHC22 | 2.094999086 | 8.125433546 | 0.621539237 | 3.370662639 | 0.000749876 |

**Table S5**. Differential expression of the COX modeled genes in *Nrf1α^−/−^* vs *WT* cell lines

| **Gene ID** | **Symbol** | ***WT*** | ***Nrf1α^−/−^*** | **log_2_(*Nrf1α^−/−^/ WT*)** | **Change** | **Probability** |
| --- | --- | --- | --- | --- | --- | --- |
| 84733 | CBX2 | 27.48 | 28.15 | 0.03 | Up | 0.20 |
| 729447 | GAGE2A | 0.01 | 0.01 | 0.00 | none | NA |
| 3235 | HOXD9 | 5.50 | 6.86 | 0.32 | Up | 0.94 |
| 6696 | SPP1 | 0.01 | 0.05 | 2.32 | Up | NA |
| 8614 | **STC2** | 3.85 | 36.97 | 3.26 | Up | 1.00 |
| 8277 | TKTL1 | 0.02 | 0.01 | -1.00 | Down | NA |

**Table S6.** Differential expression of the COX modeled genes in *Nrf2^−/−^* vs *WT* cell lines

| **Gene ID** | **Symbol** | ***WT*** | ***Nrf2^−/−^*** | **log_2_(*Nrf2^−/−^/ WT*)** | **Change** | **Probability** |
| --- | --- | --- | --- | --- | --- | --- |
| 84733 | CBX2 | 27.48 | 23.67 | -0.22 | Down | 0.96 |
| 729447 | GAGE2A | 0.01 | 0.01 | 0.00 | none | NA |
| 3235 | HOXD9 | 5.50 | 5.28 | -0.06 | Down | 0.05 |
| 8614 | **STC2** | 3.85 | 1.62 | -1.25 | Down | 1.00 |
| 8277 | TKTL1 | 0.02 | 0.01 | -1.00 | Down | NA |

**Table S7.** Differential expression of the COX modeled genes in *caNrf2^ΔN^* vs *WT* cell lines

| **GeneID** | **Symbol** | ***WT*** | *caNrf2^ΔN^* | **log_2_(ca*Nrf2****^ΔN^****/ WT*)** | **Change** | **Probability** |
| --- | --- | --- | --- | --- | --- | --- |
| 84733 | CBX2 | 27.48 | 13.54 | -1.02 | Down | 1.00 |
| 729447 | GAGE2A | 0.01 | 0.01 | 0.00 | - | NA |
| 3235 | HOXD9 | 5.50 | 10.50 | 0.93 | Up | 1.00 |
| 6696 | SPP1 | 0.01 | 0.27 | 4.77 | Up | NA |
| 8614 | **STC2** | 3.85 | 21.25 | 2.46 | Up | 1.00 |
| 8277 | TKTL1 | 0.02 | 0.01 | -1.00 | Down | NA |
| 283576 | ZDHHC22 | 0.01 | 0.06 | 2.66 | Up | NA |

**Table S8**. Distinct expression of key 46 DEGs in six examined cell lines

| **Gene *WT Nrf1α^−/−^ Nrf2^−/−^ caNrf2^ΔN^* Lentiv-STC2 *STC2^−/−^*** | | | | | | | |
| --- | --- | --- | --- | --- | --- | --- | --- |
| ACKR3 | 8.72 | 7.2 | 5.76 | | 42.46 | 3.79 | 7.63 |
| ACSL5 | 5.49 | 0.23 | 3.7 | | 12.02 | 1.63 | 10.87 |
| ADAMTSL4 | 13.39 | 6.54 | 18.62 | | 17.14 | 5.22 | 11.58 |
| ANGPTL4 | 1.06 | 10.99 | 0.69 | | 19.13 | 3.08 | 11.7 |
| BMP2 | 62.59 | 14.98 | 51.31 | | 39.28 | 27.67 | 60.39 |
| CCND1 | 14.19 | 57.37 | 25.53 | | 31.1 | 2.42 | 11.7 |
| CDH2 | 5.7 | 21.15 | 7.23 | | 12.08 | 33.34 | 0.45 |
| COL15A1 | 26.52 | 10.56 | 12.33 | | 21.26 | 7.84 | 35.64 |
| COL4A2 | 15.72 | 99.3 | 23.49 | | 53.21 | 1.57 | 14.55 |
| CSPG4 | 5.49 | 3.88 | 5.4 | | 9.12 | 3.05 | 11.24 |
| CTSF | 2.79 | 1.92 | 3.53 | | 6.67 | 1.22 | 2.36 |
| DDIT4 | 147.12 | 237.63 | 173.44 | | 250.25 | 175.01 | 334.98 |
| DUSP5 | 5.37 | 11.07 | 3.36 | | 17.98 | 9.51 | 23.54 |
| DUSP9 | 4.9 | 0.37 | 8.18 | | 3.51 | 2.38 | 6.58 |
| ECI2 | 15.86 | 12.4 | 17.77 | | 13.57 | 15.18 | 0.24 |
| ENO2 | 19.89 | 13.63 | 14.47 | | 35.52 | 19.62 | 41.01 |
| ENO3 | 35.39 | 8.96 | 33.86 | | 50.84 | 16.08 | 34.52 |
| FABP3 | 2.3 | 2.24 | 6.55 | | 10.59 | 1.19 | 3.97 |
| FGFR4 | 17.28 | 7.84 | 12.37 | | 10.57 | 13.49 | 7.72 |
| FN1 | 0.15 | 8.53 | 0.4 | | 1.18 | 0.09 | 0.97 |
| GSTO2 | 0.88 | 8.35 | 0.68 | | 5.54 | 0.54 | 2.99 |
| HEY1 | 1.27 | 1.46 | 1.85 | | 4.32 | 1.44 | 2.77 |
| HIF1A | 21.7 | 16.75 | 32.28 | | 65.74 | 28.36 | 30.75 |
| HIST2H2AC | 2.14 | 1.01 | 1.8 | | 0.69 | 1.01 | 1.85 |
| HK1 | 10.66 | 64.89 | 14.5 | | 61.76 | 2.7 | 17.14 |
| JUN | 3.34 | 17.76 | 4.38 | | 15.13 | 3.36 | 6.87 |
| KEAP1 | 54.17 | 55.93 | 45.94 | | 43.42 | 48.37 | 46.03 |
| LAMA1 | 17.08 | 6.28 | 39.82 | | 3.8 | 13.94 | 28.31 |
| LOXL2 | 30.45 | 85.27 | 13.38 | | 93.45 | 44.46 | 70.53 |
| MFSD12 | 485.4 | 266.7 | 386.97 | | 239.16 | 196.08 | 401.89 |
| MGLL | 12.91 | 17.09 | 10.7 | | 37.34 | 0.78 | 9.97 |
| MYH10 | 4.56 | 3.27 | 7.61 | | 7.63 | 0.42 | 13.61 |
| NFE2L1 | 58.92 | 27.91 | 44.97 | | 135.43 | 70.45 | 57.11 |
| NFE2L2 | 14.83 | 11.39 | 19.18 | | 25.77 | 15.71 | 14.76 |
| NGEF | 3.81 | 6.4 | 2.46 | | 0.33 | 0.52 | 4.22 |
| NR4A1 | 15.9 | 24.56 | 14.66 | | 19.27 | 10.55 | 22.38 |
| NRP2 | 1.91 | 5.63 | 0.71 | | 0.85 | 0.61 | 2.19 |
| PCDH7 | 5.97 | 5.78 | 11.69 | | 6.96 | 53.49 | 1.03 |
| PTGS1 | 2.93 | 0.04 | | 0.77 | 0.04 | 2.94 | 0.92 |
| SOHLH2 | 0.16 | 0.06 | | 0.04 | 2.1 | 0.00 | 0.36 |
| SPOCK1 | 0.61 | 4.28 | | 1.69 | 5.33 | 1.95 | 0.27 |
| SQSTM1 | 849.41 | 608.53 | | 420.76 | 1065.43 | 644.8 | 331.52 |
| STC2 | 3.85 | 36.97 | | 1.62 | 21.25 | 126.69 | 0.33 |
| TFEB | 2.94 | 7.13 | | 2.6 | 2.92 | 3.97 | 2.64 |
| TGFB3 | 1.2 | 3.24 | | 1.55 | 0.71 | 0.99 | 2.35 |
| TNNC1 | 49.89 | 0.85 | | 69.57 | 6.49 | 21.39 | 34.84 |
